# High performance machine learning models can fully automate labeling of camera trap images for ecological analyses

**DOI:** 10.1101/2020.09.12.294538

**Authors:** Robin Whytock, Jędrzej Świeżewski, Joeri A. Zwerts, Tadeusz Bara-Słupski, Aurélie Flore Koumba Pambo, Marek Rogala, Laila Bahaa-el-din, Kelly Boekee, Stephanie Brittain, Anabelle W. Cardoso, Philipp Henschel, David Lehmann, Brice Momboua, Cisquet Kiebou Opepa, Christopher Orbell, Ross T. Pitman, Hugh S. Robinson, Katharine A. Abernethy

## Abstract

1. Ecological data are increasingly collected over vast geographic areas using arrays of digital sensors. Camera trap arrays have become the ‘gold standard’ method for surveying many terrestrial mammals and birds, but these arrays often generate millions of images that are challenging to process. This causes significant latency between data collection and subsequent inference, which can impede conservation at a time of ecological crisis. Machine learning algorithms have been developed to improve camera trap data processing speeds, but these models are not considered accurate enough for fully automated labeling of images.
2. Here, we present a new approach to building and testing a high performance machine learning model for fully automated labeling of camera trap images. As a case-study, the model classifies 26 Central African forest mammal and bird species (or groups). The model was trained on a relatively small dataset (*c*.300,000 images) but generalizes to fully independent data and outperforms humans in several respects (e.g. detecting ‘invisible’ animals). We show how the model’s precision and accuracy can be evaluated in an ecological modeling context by comparing species richness, activity patterns (*n* = 4 species tested) and occupancy (*n =* 4 species tested) derived from machine learning labels with the same estimates derived from expert labels.
3. Results show that fully automated labels can be equivalent to expert labels when calculating species richness, activity patterns (*n* = 4 species tested) and estimating occupancy (*n* = 3 of 4 species tested) in completely out-of-sample test data (*n* = 227 camera stations, *n* = 23868 images). Simple thresholding (discarding uncertain labels) improved the model’s performance when calculating activity patterns and estimating occupancy, but did not improve estimates of species richness.
4. We provide the user-community with a multi-platform, multi-language user interface for running the model offline, and conclude that high performance machine learning models can fully automate labeling of camera trap data.

## Introduction

The urgent need to understand how ecosystems are responding to rapid environmental change has driven a ‘big data’ revolution in ecology and conservation (Farley, Dawson, Goring, & Williams, 2018). High resolution ecological data are now streamed in real-time from satellites, Global Positioning System tags, bioacoustic detectors, cameras and other sensor arrays. The data generated offer considerable opportunities to ecologists, but challenges such as data processing, data storage and data sharing cause latency between data gathering and ecological inference (i.e. creating derived ecological metrics, testing ecological hypotheses and quantifying ecological change), sometimes in the order of years or more. Overcoming these challenges could open the gateway to ecological ‘forecasting’, where directional changes in ecological processes are detected in real time and near-term responses are predicted effectively using an iterative data gathering, model updating and model prediction approach (Dietze et al., 2018).

Digital camera traps or wildlife ‘trail cams’ have revolutionized wildlife monitoring and are now the ‘gold standard’ for monitoring many medium to large terrestrial mammals (Glover-Kapfer, Soto‐ Navarro, & Wearn, 2019). Animals and their behavior are identified in images either by manual labeling, using citizen science platforms (Swanson et al., 2015) or, more recently, by using machine learning models (Norouzzadeh et al., 2018; Tabak et al., 2019; Willi et al., 2019). Machine learning models can at minimum separate true animal detections from non-detections (Wei, Luo, Ran, & Li, 2020) or in the most advanced examples identify species, count individuals and describe behavior (Norouzzadeh et al., 2018). These recent advances in machine learning have increased the speed at which camera trap data are analyzed but, in all cases we are aware of, the outputs (e.g. species labels) are not used to make ecological inference directly. Instead, machine learning models are typically used as a ‘first pass’ to identify and group images belonging to individual species for full or partial manual validation at a later stage, or to cross-validate labels from citizen science platforms (Willi et al., 2019). This can substantially reduce manual labeling effort but many hundreds or thousands of photos might still need to be labeled manually. Thus, although machine learning models are reducing manual data processing times, ecologists are not yet comfortable using the outputs (e.g. species labels) as part of a completely automated workflow. This is despite the development of advanced machine learning models that classify species in camera trap images with accuracy that matches or exceeds humans (Norouzzadeh et al., 2018; Tabak et al., 2019).

One significant challenge limiting the application of machine learning models to camera trap data is that models rarely generalize well to completely out-of-sample data (i.e. data from new, spatially and temporally independent studies), particularly when used to classify animals to species level (Beery, Van Horn, & Perona, 2018). Models can quickly learn the features of specific camera ‘stations’ (the spatial replicate in camera trap studies) such as the general background instead of learning features of the animal itself. This problem is further amplified by the fact that rare species in the training data might only ever appear at a limited number of camera stations, so training and validation data are rarely independent. Various approaches can be used to reduce these biases, such as carefully ensuring that training and validation data are independent (e.g. by using data from multiple studies), and by using data augmentation such as adding noise to training data in the form of image transformations. Until the problem of generalization can be overcome, machine learning models for classifying camera trap images will remain an important tool for reducing manual labeling effort, but they will not achieve their full potential for creating fully automated pipelines for data analysis.

Machine learning models also have the potential to be deployed inside camera trap hardware in the field at the ‘edge’ (i.e. on micro-computers installed inside hardware that collects data), with summarized results (e.g. species labels) transmitted in real-time via a Global System for Mobile Communications networks or via satellite (Glover-Kapfer et al., 2019). In geographically remote areas or time-sensitive situations (e.g. law enforcement) this would greatly reduce the latency between data capture and interpretation, and reduce the expense and effort required to collect data in remote regions by removing the need to transfer data-heavy images across wireless networks. However, before ‘smart’ cameras become a reality, it is essential that users understand how uncertainty in machine learning model predictions might impact derived ecological metrics and analyses, which are often sensitive to biases (e.g. false positives in occupancy models). To achieve this, there is a need to develop workflows that test the performance of machine learning models in an ecological modeling context that goes beyond simple measures of precision and accuracy.

Ideally, if machine learning models had 100% precision and accuracy (e.g. for species identification), camera trap data could be collected, labeled automatically using the model and the results used to directly calculate ecological metrics or as variables in ecological models. However, the reality is that machine learning models are imperfect. It is therefore uncertain what levels of precision and accuracy are needed to meet the requirements of ecological analyses. This is particularly the case for the spatial and temporal analayses of animal distributions in camera trap data, which require specialized ecological models (e.g. occupancy models) that account for imperfect detection (MacKenzie et al., 2002).

In this paper, we describe the approach used to build a new high-performance machine learning model that identifies species in camera trap images (26 species/groups of Central African forest mammals and birds) that generalizes to spatially independent data. To evaluate how well the machine learning model labeling precision and accuracy performs in an ecological modeling context, we (1) evaluate how uncertainties in the precision and accuracy of machine learning labels affect ecological inference (derived metrics of species richness, activity patterns and occupancy) compared to the same metrics calculated using expert, manually generated labels, and (2) propose a workflow to ‘ground truth’ the performance of machine learning models for camera trap data in an ecological modeling context. We discuss the implications of these results for making fully automated ecological inference from camera trap data using the outputs of machine learning models. We also provide the user community with an easily installed, open-source graphical user interface that needs no understanding of machine learning to run the model offline on both camera trap images and videos.

## Methods

### Data preparation

As a case study, the model was developed for classifying terrestrial forest mammals and birds in Central Africa (see Table S1 for further details on species and groups), where camera traps are now frequently deployed over large spatial scales to survey secretive birds and mammals in remote and inaccessible landscapes (Bahaa-el-din & Cusack, 2018; Bessone et al., 2020; O’Brien et al., 2020). Training data were obtained from multiple countries and sources (*c*.1.6 million images; reduced to *n* = 347120 images after data processing; Table 1). Each source used different camera trap models (Reconyx, Bushnell, Cuddeback, Panthera Cams) and images were diverse in resolution, quality (e.g. sharpness, illumination) and color. Individual studies also used different field protocols for camera deployment but all were focused on detecting terrestrial forest mammals, with cameras installed on trees approximately 30 - 40 cm above ground level. The exception to this was data from (Cardoso et al., 2020) who installed cameras at a height of approximately 1 m for the primary purpose of detecting forest elephants *Loxodonta cyclotis*. Camera trap configuration was set to be highly sensitive in some cases and images were often captured in a series of rapid, short bursts (e.g. taking 10 images consecutively). This resulted in long sequences of very similar images, for example showing an animal walking in front of the camera (Figure S1).

**Table 1.** Sources of training data used to train the machine learning model for classifying species in camera trap images, sorted by number of images provided. The final subset of data used to train the model was *n* = 347120 images (see later).

| Source | Country | Reference | *n* images |
| --- | --- | --- | --- |
| Anabelle Cardoso | Gabon | (Cardoso et al., 2020) | 102418 |
| Kelly Boekee | Cameroon | - | 123954 |
| Cisquet Kiebou Opepa | Republic of Congo | - | 60393 |
| Joeri Zwerts | Cameroon | - | 36027 |
| Laila Bahaa-el-Din | Gabon | (Bahaa-el-din et al., 2013) | 16558 |
| Stephanie Brittain | Cameroon | - | 7770 |

It was important to account for image sequences when selecting a validation set during the model training phase, since there was a risk of highly similar images being present in both the training and validation sets. To address this issue, the training and validation split was performed based on image metadata (timing of images and image source) to identify unique ‘events’ and camera locations that were not replicated across the training and validation split (Norouzzadeh et al., 2018). This solution posed a challenge for maintaining class balances in the training and validation sets, but it reduced the risk nonindependent training and validation sets. A total of 27 classes were used to train the model, which were mostly mammals or mammal groups (*n* = 21), birds (*n* = 4), humans (*n =* 1) and ‘blank’ images (i.e. no mammal, bird or human). Details of taxonomy and justification for species groups are in Table S1.

### Issues identified in the training data

Our ‘real-life’ training data had not been pre-processed or professionally curated for the purposes of training machine learning models and naturally contained errors that arise from hardware faults, human error and different approaches to manual species labeling by experts. We identified three primary sources of error. The first was over-exposed images (a hardware fault) where the image foreground was ‘flooded’ by the flash (usually at night), making the image appear mostly white. Animals in these images were sometimes partially visible and could be classified by a skilled human observer, despite the loss of color information, texture and other detail. However, over-exposed images presented a challenge for the machine learning model because white dominated the image regardless of the species.

The second main source of error was caused by under-exposed images. This error was revealed after inspecting model outputs during the training phase, and showed that highly under-exposed images appeared almost entirely or entirely black to a human observer, but the machine learning model was capable of using information in the image to detect and correctly classify the species (Figure 1).

**Figure 1.**
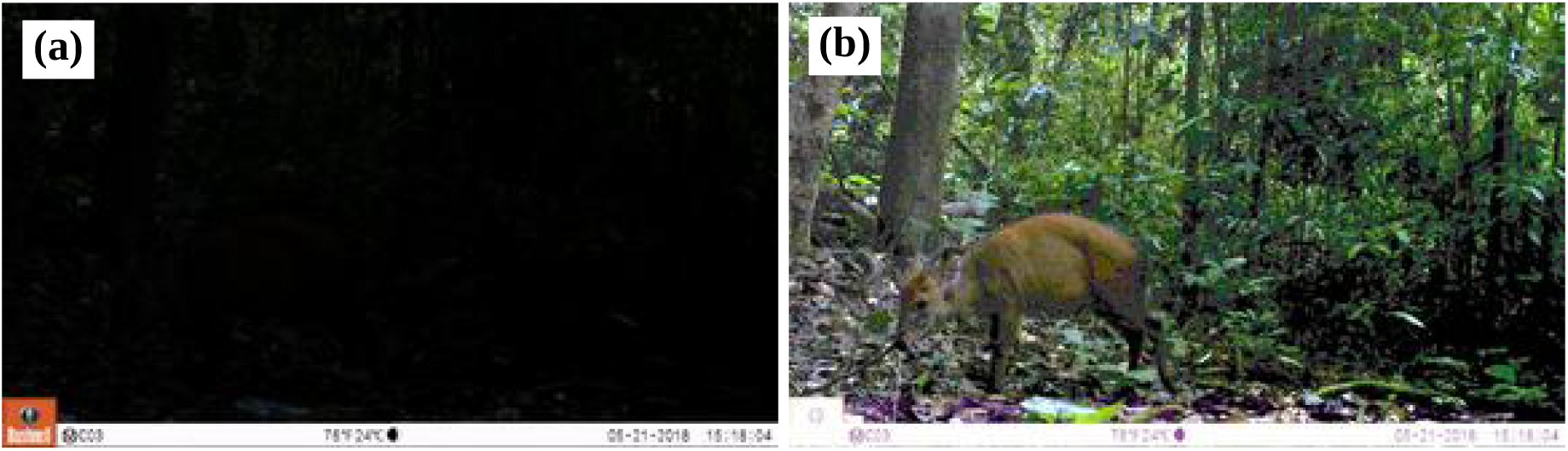
(a) Raw image from the dataset, labeled by experts as “blank”, but classified by the machine learning model with high certainty as a red duiker. (b) The same image as in (a), but manually brightened by narrowing the displayed color spectrum, reveals a red duiker is present and the model was correct.

The final source of error in the training data was mis-labeled images (e.g. confusing similar species, such as chimpanzee *Pan troglodytes* and gorilla *Gorilla gorilla*) and using different approaches to labeling, for example one data source combined all primates into ‘monkey’, whereas other data sources separated apes from other primates.

We used an iterative approach to address these issues that consisted of model training, validation, error correction (correcting mis-labeled images in the training data) and model updating. In particular, we carefully inspected images that appeared to be incorrectly labeled by the model, but which were labeled with high confidence. This approach revealed hidden problems in the data, such as the presence of animals in under-exposed images that would have otherwise led us to underestimate the model’s performance.

### Machine learning model

We chose the established ResNet50 architecture to build the model (He, Zhang, Ren, & Sun, 2016). Transfer learning was used to speed up training and we used weights pre-trained on the ImageNet dataset. We identified species using the entire image frame without using bounding boxes and used basic augmentation (horizontal flips, rotations, zoom, lighting and contrast adjustments, and warps) during training, but not during model validation. We used one-cycle policy training (Smith, 2018) and trained using progressive resizing in two stages. Details on the training scheme and implementation can be found in our GitHub repository (https://github.com/Appsilon/gabon_wildlife_training). It is worth noting that most of the training approaches and many of the mechanisms we used to enhance training were taken directly or almost directly from the fast.ai Python library (https://github.com/fastai), exemplifying how exceptionally robust the library is. We trained the models on various virtual machines equipped with GPU processing units, run on Google Cloud Platform with resources granted by a Google Cloud Education grant.

### Out-of-sample test data

One of the major limitations to model performance for camera trap images is the ability to generalize predictions to new, independent camera stations, i.e. unique locations with different backgrounds not seen during model training (Beery et al., 2018). Since our objective was to create a model that could generalize well to new study sites, we tested the final model’s performance using a new out-of-sample dataset that was completely spatially and temporally independent from the data used to train the model. These out-of-sample data consisted of images from 227 camera stations surveyed between 16 January 2018 and 4 October 2019 in central and southern Gabon in closed canopy forest. Cameras also differed from the models used in the training data (Panthera Cams V4 and V5), but field protocols were similar and cameras were placed approximately 30 cm above the ground on a tree at a distance of c. 3 – 5 m perpendicular to the center of animal trails. Single-frame images were captured using medium sensitivity settings, and images were separated by a minimum of 1 s. The aim of the study was to survey the small-to-large mammal community, with a particular focus on great apes (*Pan troglodytes, Gorilla gorilla*), forest elephants *Loxodonta cyclotis*, leopard *Panthera pardus* and golden cat *Caracal aurata*. These data (*n* = 23868 images, median 75, range 1 - 545 images per station) were manually labeled by an expert (co-author CO).

### Summary of model’s general performance

To allow general comparison of our model’s performance with other similar models in the literature (Norouzzadeh et al., 2018; Tabak et al., 2019; Willi et al., 2019) we calculated top-one and top-five accuracies using the out-of-sample data. Top-one accuracy is the percent of expert labels that match the top-ranking label generated by the machine learning model. Top-five accuracy calculates the percent of expert labels that match any of the top five ranking machine learning generated labels. Top-one accuracy for the overall machine learning model was 77.63% and top-five accuracy was 94.24% (Table S2; Figures S2 & S3). After aggregating labels of similar species that were frequently mis-classified by the model into a reduced set of 11 classes, top-one and top-five accuracies increased to 79.92% and 95.99%, respectively (Figure S4). The model can classify around 4000 images (c.0.5 MB in size) per hour using an Intel^®^ Core™ i7-8665U CPU @ 1.90GHz × 8 and the model can operate 24/7 if necessary. For comparison, based on our experience, manual labeling can be done at speeds ranging from 125 to 500 images per hour depending on the quality of the images and if images are captured in sequences (which can be faster to label manually).

We also compared the precision and recall for each species from our optimal model (see later, Table 2) with precision and recall for the same species reported for the model used by the WildlifeInsights webplatform (www.wildlifeinsights.org). This global project uses a deep convolutional neural network trained using Google’s Tensorflow framework and a training dataset of 8.7M images, comprising 614 species.

### Comparing derived ecological metrics using machine learning labels and expert labels

We calculated three common ecological metrics for the out-of-sample data (raw species richness at individual camera stations, activity patterns for four focal species, and occupancy for four focal species) separately using the manually generated, expert labels and the machine learning generated labels. Species richness (the number of species in a discrete unit of space and time) can be used to quantify temporal and spatial changes in biodiversity. Although other measures of species diversity exist, we chose this simple metric because it is widely used in the ecology literature despite its limitations. Activity patterns describe the diel activity patterns of focal species (M. Rowcliffe, 2019) and are typically calculated to understand fundamental life history traits and behavior such as temporal niche partitioning. Occupancy models are hierarchical models commonly fitted to camera trap data because they can account for imperfect detection (which rarely equals 1) to estimate the conditional probability that a site is ‘occupied’ by a species given it was not detected (MacKenzie et al., 2002). Covariates such as measures of vegetation cover can be included in both the detection and occupancy component models. These models are relatively complex, and small changes in detection histories (presence or absence of a species during a discrete time interval), false positives or false negatives can dramatically affect results (Royle & Link, 2006). We therefore predicted that occupancy estimates obtained using machine learning generated labels would compare poorly with estimates using expert, manually generated labels.

The four focal species used for calculating activity patterns and occupancy were African golden cat, chimpanzee, leopard and African forest elephant. These species were chosen because they were the focus of the camera trap survey that generated the out-of-sample test data and because they are conservation priority species in Central Africa. We also initially included western lowland gorilla but we had too few unique captures of this species (only seven of 227 stations having > 5 captures) to fit either activity pattern models or occupancy models.

### Thresholding and overall model performance

All three metrics derived from machine learning labels were re-calculated using a threshold approach, where labels were excluded if the model’s predicted confidence was below a given threshold. The thresholds tested ranged from 0 (no threshold) to 90%, increasing in 10% intervals. For each of the three ecological metrics, we then re-calculated results using the machine learning labels and compared these with results from the expert labeled dataset using various statistical measures (see later). We also calculated the effect of removing data on sample size, top-one balanced accuracy and top-five accuracy for the overall model, and on four standard measures of model precision and accuracy (precision, recall, F1 score, and balanced accuracy for each species using the confusionMatrix function in the caret R package (Kuhn, 2020).

Estimated species richness from machine learning generated labels and expert labels was compared using linear regression fitted by least squares. Species richness from expert labels was used as the predictor variable and species richness from machine learning labels was used as the response. For each threshold, we evaluated how well species richness from machine learning labels correlated with expert labels by calculating the slope coefficient and variance explained (R^2^).

Diel activity patterns were calculated for all four focal species using the fitact function (200 bootstrap replicates from the model) using the activity R package (J. M. Rowcliffe, Kays, Kranstauber, Carbone, & Jansen, 2014; M. Rowcliffe, 2019). For each species and threshold combination, we tested if there was a significant difference in diel activity (proportion of 24 h day active) estimated by machine learning labels and expert labels using the compareAct function, expecting no difference using an alpha level of 0.05.

Single season, single species occupancy models were fitted using the occu function from the unmarked R package (Fiske & Chandler, 2011). Detection histories were collapsed to five-day occasion lengths as a compromise between achieving model stability and ensuring an adequate number of replicates for each site. In the detection component model, we included Elevation (m), Date (first day of the five day occasion length) and Date^2^ (to allow for non-linear, seasonal changes in detection) as covariates. In the occupancy component model, Elevation (m), Distance to the Nearest River (m), Distance to the Nearest Road (m) and mean distance to the Nearest Village (m) were included as continuous predictors without interactions. All covariates were mean-centered and scaled by 1 SD to prevent convergence issues. We did not perform model selection and predicted occupancy for the 227 camera stations using the full model. We then compared occupancy predictions (*n* = 227 camera stations) for no threshold (i.e. using all data), and the nine thresholds using linear regression fitted by least squares as described previously for the species richness comparison.

## Results

### Effect of thresholding on overall model performance

Regardless of the threshold used, top-five accuracy for the overall model predictions on the out-ofsample data were consistently close to or above 95% (Figure 2). To achieve a top-one balanced accuracy of 90% or more for the overall model, a threshold of ≥ 70% confidence was required and > 25% of the data were discarded (Figure 2). With a threshold of 70% confidence (i.e. excluding labeled images below 70% confidence), top-one balanced accuracies for 16 of the 27 classes were > 90% and a further five were > 75% (Table 2). Top-one balanced accuracies for the remaining seven classes ranged from 50% to 70% (Table 2). All other measures of accuracy and precision at all thresholds are in Table S3 and Figure 3 shows the confusion matrix for the out-of-sample data after excluding labels below 70% confidence (see Figure S5 for the confusion matrix of aggregated labels after thresholding).

**Figure 2.**
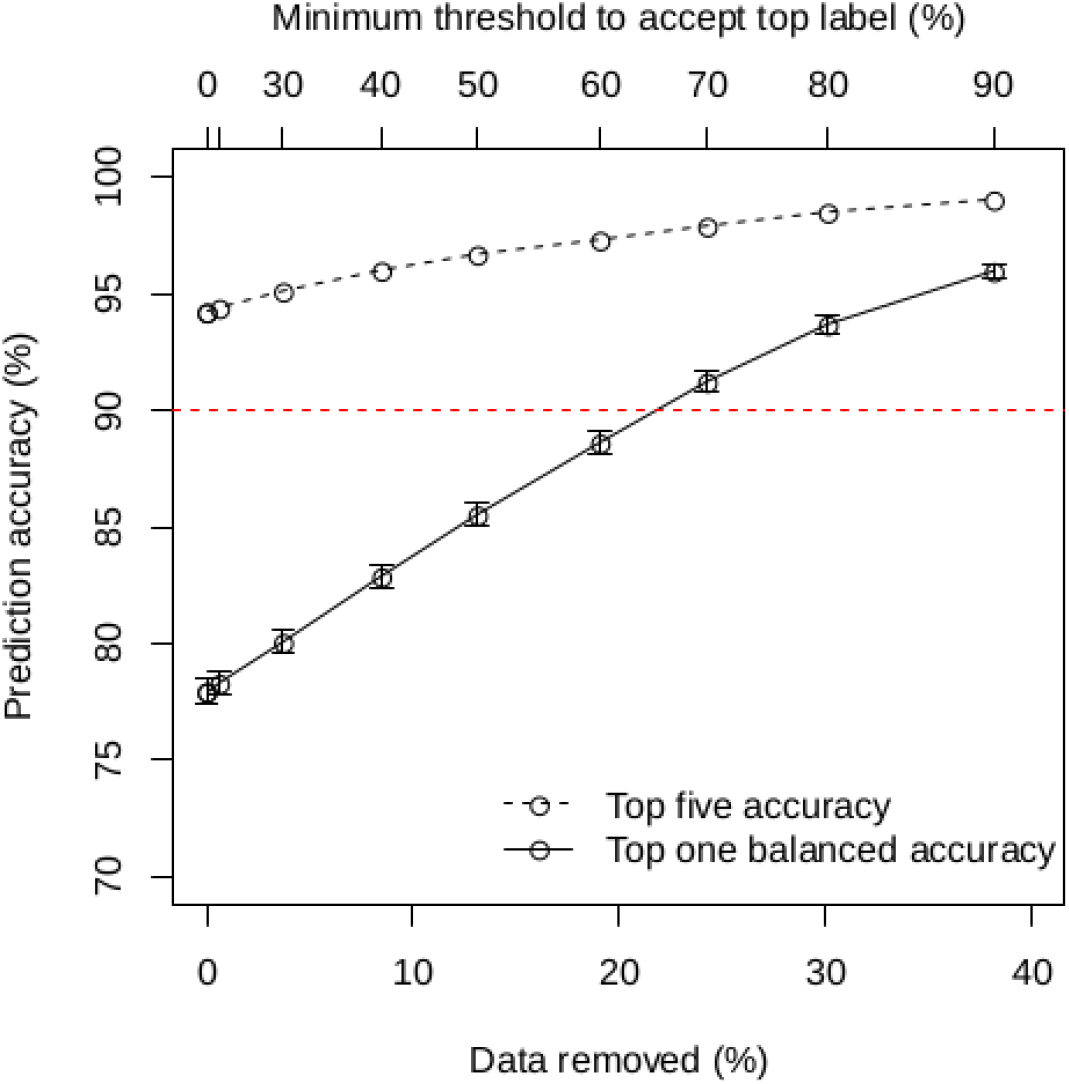
Relationship between threshold level to accept top label, % of data discarded and overall top-five and top-one balanced accuracy (+/- 95% CI) for predictions on out-of-sample test data.

**Figure 3.**
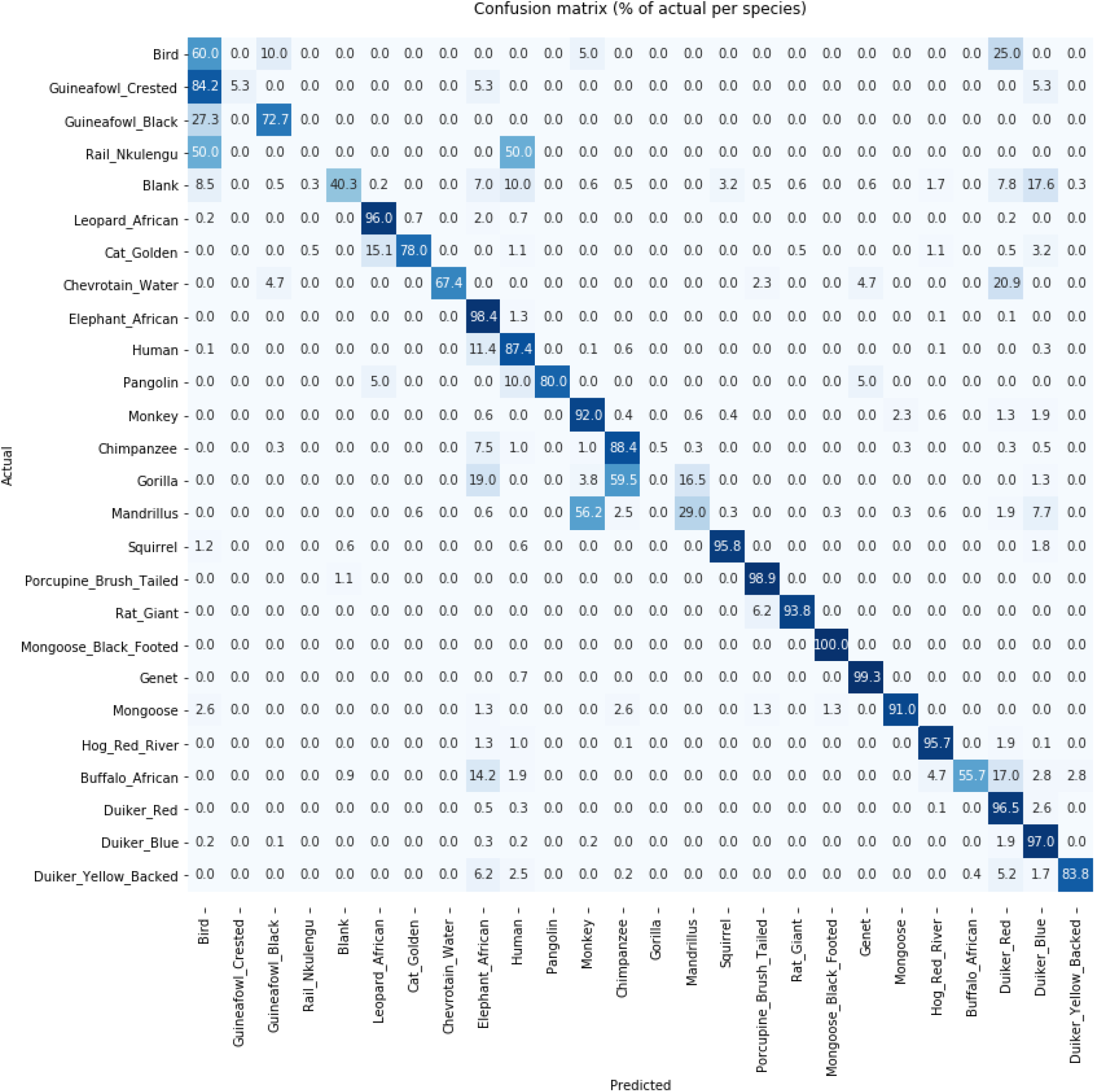
Confusion matrix (% correct labels for each species/group) showing model performance on out of sample test data after excluding labels below a confidence threshold of 70% (each row is normalized independently). Figure S6 shows the confusion matrix with absolute numbers.

**Table 2.** Precision, recall, accuracy, F1 score and prevalence (%s) for the 27 classes (Table S1) in the out-of-sample test data after removing labels with a predicted confidence < 70%. Species are sorted from lowest to highest balanced accuracy. For comparison, the precision and recall for the model used by the wildlifeinsights.org web platform are given in brackets. Orange indicates our model performed worse than the WildlifeInsights model for a given species, and purple indicates our model performed better. Note that this comparison should be interpreted with caution. Ideally, we would run the WildlifeInsights model on our out-of-sample test data, but data sharing restrictions prevented this. Where our species or groups could not be compared with an equivalent class on WildlifeInsights this is indicated as no equivalent class (NE). If precision and recall cannot be estimated because of insufficient training and validation data this is indicated as ‘needs more data’ (NMD).

| Species | Precision % | Recall % | F1 | Prevalence | Balanced Accuracy |
| --- | --- | --- | --- | --- | --- |
| Civet_African_Palm | NMD (*NMD*) | NMD (*NMD*) | NA | NA | NA |
| Gorilla | NMD (*NMD*) | NMD (*NMD*) | NA | 0.4 | 50 |
| Rail_Nkulengu | **0.0** (47.2) | **0.0** (48.6) | NA | NA | 50 |
| <sup>a</sup><br>Guineafowl_Crested | 100 (99.8) | **5.3** (91.2) | 10 | 0.1 | 52.6 |
| Mandrillus | **83.9** (96.1) | **29** (72.3) | 43.1 | 1.8 | 64.5 |
| Blank | 98.1 (98.3) | **40.3** (78.7) | 57.1 | 3.6 | 70.1 |
| Buffalo_African | **97.5** (91.1) | **55.7** (73.6) | 70.9 | 1.2 | 77.8 |
| Bird | 11.2 (*NE*) | 60.0 (*NE*) | 18.9 | 0.1 | 79.7 |
| Chevrotain_Water | **100** (*NMD*) | **67.4** (*NMD*) | 80.6 | 0.2 | 83.7 |
| Guineafowl_Black | **70.6** (79.6) | **72.7** (79.5) | 71.6 | 0.2 | 86.3 |
| Cat_Golden | **96.0** (*NMD*) | **78.0** (*NMD*) | 86.1 | 1 | 89 |
| Pangolin | **94.1** (*NMD*) | **80.0** (*NMD*) | 86.5 | 0.1 | 90 |
| Duiker_Yellow_Backed | **97.5** (88.8) | **83.8** (72.3) | 90.2 | 2.9 | 91.9 |
| Human | **78.4** (84.8) | **87.4** (75.2) | 82.6 | 4 | 93.2 |
| Chimpanzee | **83.5** (87) | **88.4** (71.4) | 85.9 | 2.2 | 94 |
| Monkey | 70.7 (*NE*) | 92.0 (*NE*) | 80 | 2.9 | 95.4 |
| Mongoose | **83.5** (*NMD*) | **91.0** (*NMD*) | 87.1 | 0.4 | 95.5 |
| Rat_Giant | **68.2** (76) | **93.8** (75.8) | 78.9 | 0.1 | 96.9 |
| <sup>b</sup><br>Duiker_Red | **95.9** (95.6) | **96.5** (79.6) | 96.2 | 30.8 | 97.3 |
| Duiker_Blue | **90.04** (98.2) | **97.0** (65.7) | 93.6 | 17.6 | 97.4 |
| Hog_Red_River | **97.0** (82.7) | **95.7** (84.7) | 96.3 | 6.5 | 97.7 |
| Squirrel | **85.9** (98.6) | **95.8** (67.6) | 90.6 | 0.9 | 97.8 |
| Leopard_African | **92.8** (85.2) | **96.0** (61.4) | 94.4 | 2.2 | 97.9 |
| Elephant_African | **91.9** (94.4) | **98.4** (84.2) | 95.1 | 19.3 | 98.2 |
| Porcupine_Brush_Tailed | **93.9** (89.4) | **98.9** (42.1) | 96.3 | 0.5 | 99.4 |
| Genet | **95.3** (89.2) | **99.3** (65.6) | 97.2 | 0.8 | 99.6 |
| Mongoose_Black_Footed | **92.9** (*NMD*) | **100** (*NMD*) | 96.3 | 0.1 | 100 |

Used precision and recall for similar *Guttera plumifera* from WildlifeInsights
Used precision and recall for *Cephalophus callipygus* from WildlifeInsights

### Species richness

Species richness estimated by machine learning labels and expert labels was strongly correlated at all thresholds used (Figure 4). There was a general tendency for species richness to be underestimated by machine learning as the threshold increased, and the slope of the relationship was close to 1 with no threshold.

**Figure 4.**
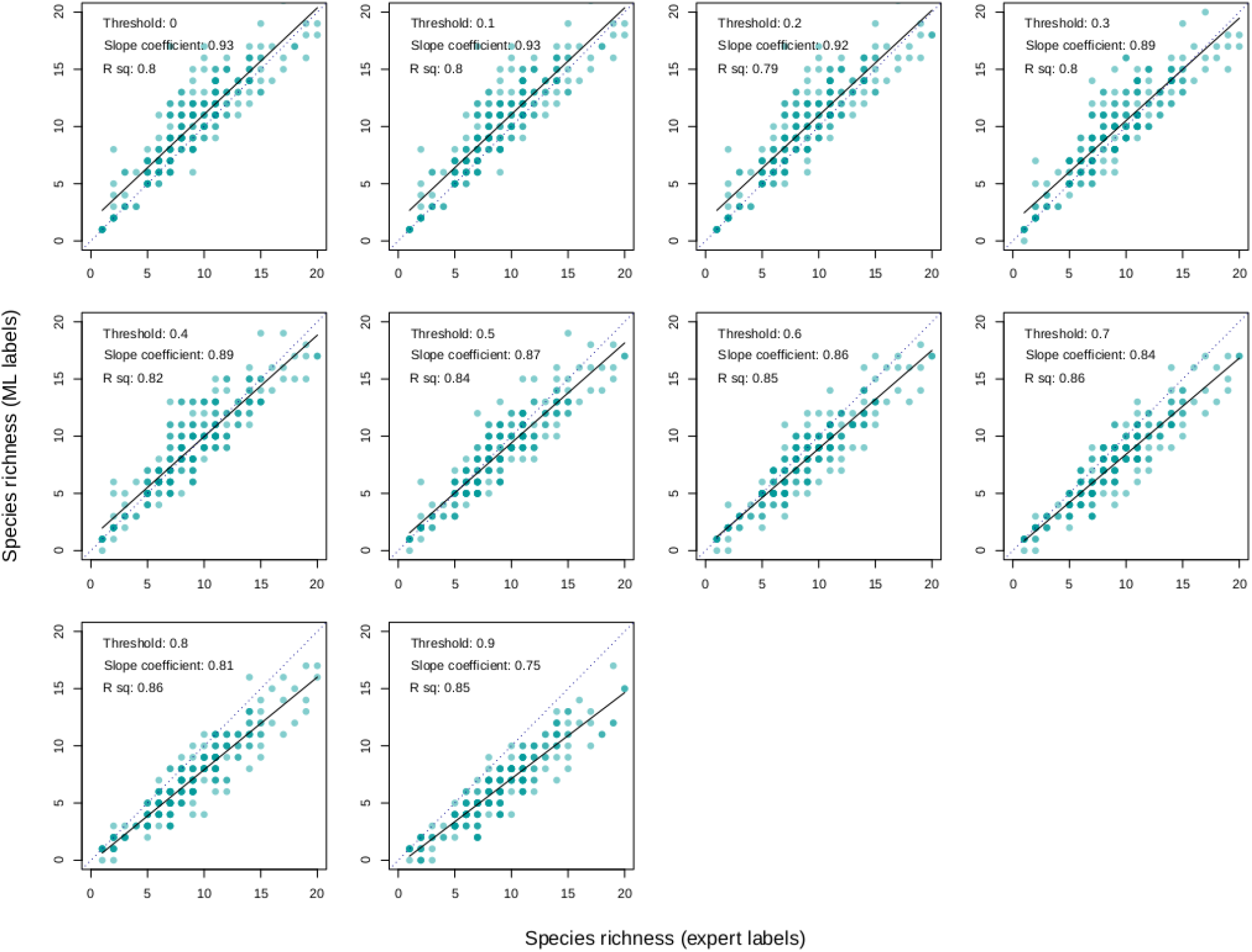
Relationship between species richness at each camera station (*n* = 227) predicted by the machine learning model (y-axis) and species richness predicted from expert labels (x-axis) for no threshold and the nine thresholds used after predicting on the out-of-sample test data. The dotted line shows where a 1:1 relationship would fit the data.

### Activity patterns

Above a threshold of 70% there was no significant difference between diel activity patterns estimated by machine learning labels and expert labels for all four focal species in the out-of-sample test data (Figure 5; Table S4).

**Figure 5.**
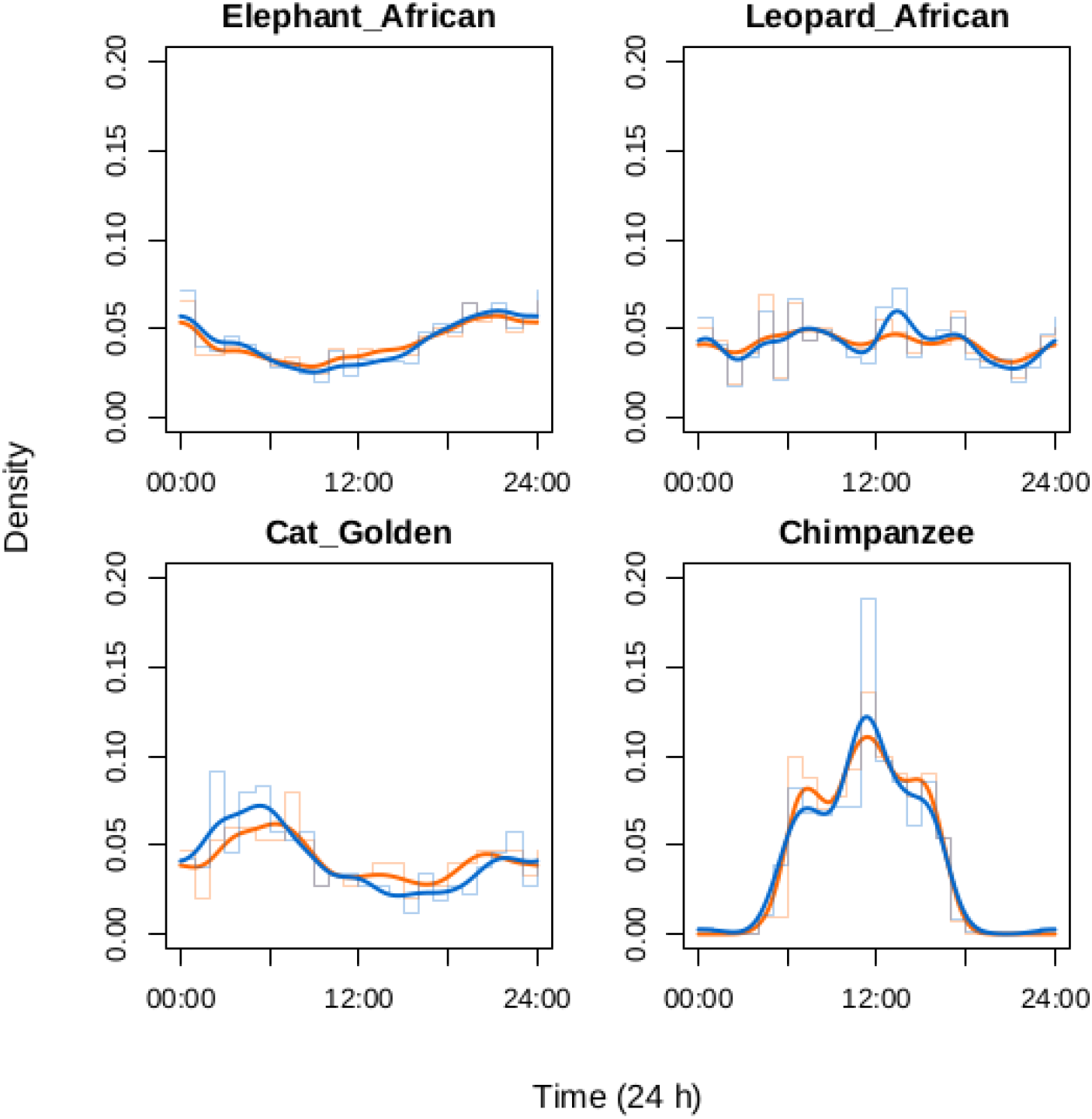
Estimated activity patterns for the four focal species in the out-of-sample test data using machine learning labels (orange; *n* = 18078 observations after excluding labels below 70% confidence) and expert labels (blue; *n* = 23868 observations).

### Occupancy models

As expected, occupancy estimates made using machine learning labels were sometimes inconsistent with those made using expert labels, and thresholding had a dramatic impact on inference in some cases (Figure 6). For golden cat and leopard, which are predicted with high accuracy and precision by our machine learning model, occupancy estimates from machine learning labels and expert labels were highly correlated at all thresholds (Figure S8). African elephant occupancy estimates using machine learning labels improved dramatically as the threshold increased, but chimpanzee occupancy estimates from machine learning labels were consistently uncorrelated with those estimated using expert labels (Figure 6).

**Figure 6.**
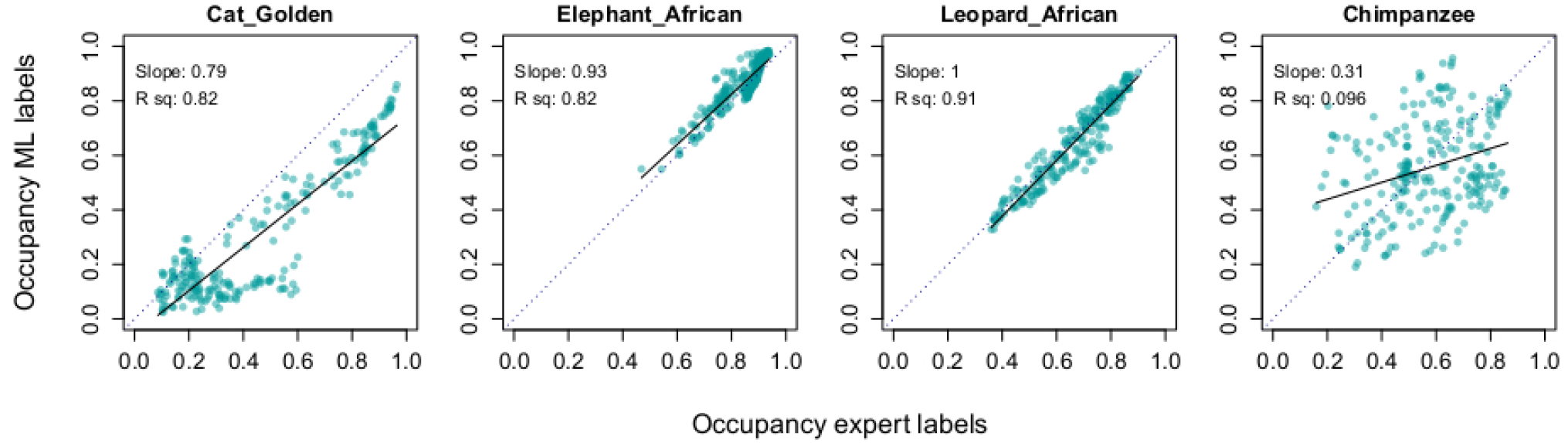
Relationship between estimated occupancy probability for *n* = 227 camera stations (points) from machine learning (ML) labels (y-axis) and expert labels (x-axis) for the four focal species after discarding labels below a 90% threshold of predicted confidence. Plots for all thresholds tested are shown in Figure S8.

## Discussion

Machine learning models have the potential to fully automate labeling of camera trap images without the need for manual validation. This would allow ecologists to rapidly process data and use the outputs (e.g. species labels) directly in ecological analyses, but it has been uncertain how this can be achieved.

In particular, models published to date do not evaluate their predictive performance in an ecological modeling context (Beery et al., 2018; Norouzzadeh et al., 2018; Tabak et al., 2019; Willi et al., 2019). Here, we compared ecological metrics calculated on an out-of-sample test dataset using machine learning labels with the same metrics calculated using expert, manually generated labels. Using our new, high performance species classification model that generalizes to out-of-sample data, we show machine learning labels can be used in a fully automated workflow that removes the need for manual validation prior to conducting ecological analyses.

We used an established architecture for the machine learning model. However, other more recent architectures could yield further increases in performance. The ResNeXt (Xie, Girshick, Dollar, Tu, & He, 2017), the ResNeSt (Zhang et al., 2020) and the EfficientNet (Tan & Le, 2020) families of network architectures are particularly worth exploring in this context. Another avenue of possible further improvement is to use an approach based on a sequence of models. One natural step is to first detect a bounding box for an animal with a localisation model (Beery et al., 2019) and later classify only the content found in that box. Independently, another step can be introduced where a model is trained to first identify an aggregated species class (comprised of species that share similar characteristics; e.g. see Figure S4), and later dedicated models are trained to identify the individual species within these aggregated classes.

We used a relatively small training set (*c*.300,000 images here *vs* 3.2 million in (Norouzzadeh et al., 2018) and 8.7M used by (Ahumada et al., 2020)) and a large number of individual classes, yet our model achieved high precision and accuracy even when tested on completely out-of-sample data, which is considered a significant challenge for the field (Beery et al., 2018, 2019). We believe this encouraging result can be explained both by the machine learning approaches used (e.g. the fast.ai framework and image augmentation), and because forest camera traps in the tropics are often deployed in very similar settings, with animals captured at a predictable distance from the camera (usually on a path) with a general background of green and brown vegetation. This is in contrast to camera trap images from more open habitats, where animals are detected across a wide range of distances and backgrounds (Beery et al., 2018). On the other hand, informational richness in the background of photos taken in forest settings poses a significant challenge to machine learning models as well as human experts, as illustrated in Figure 7.

**Figure 7.**
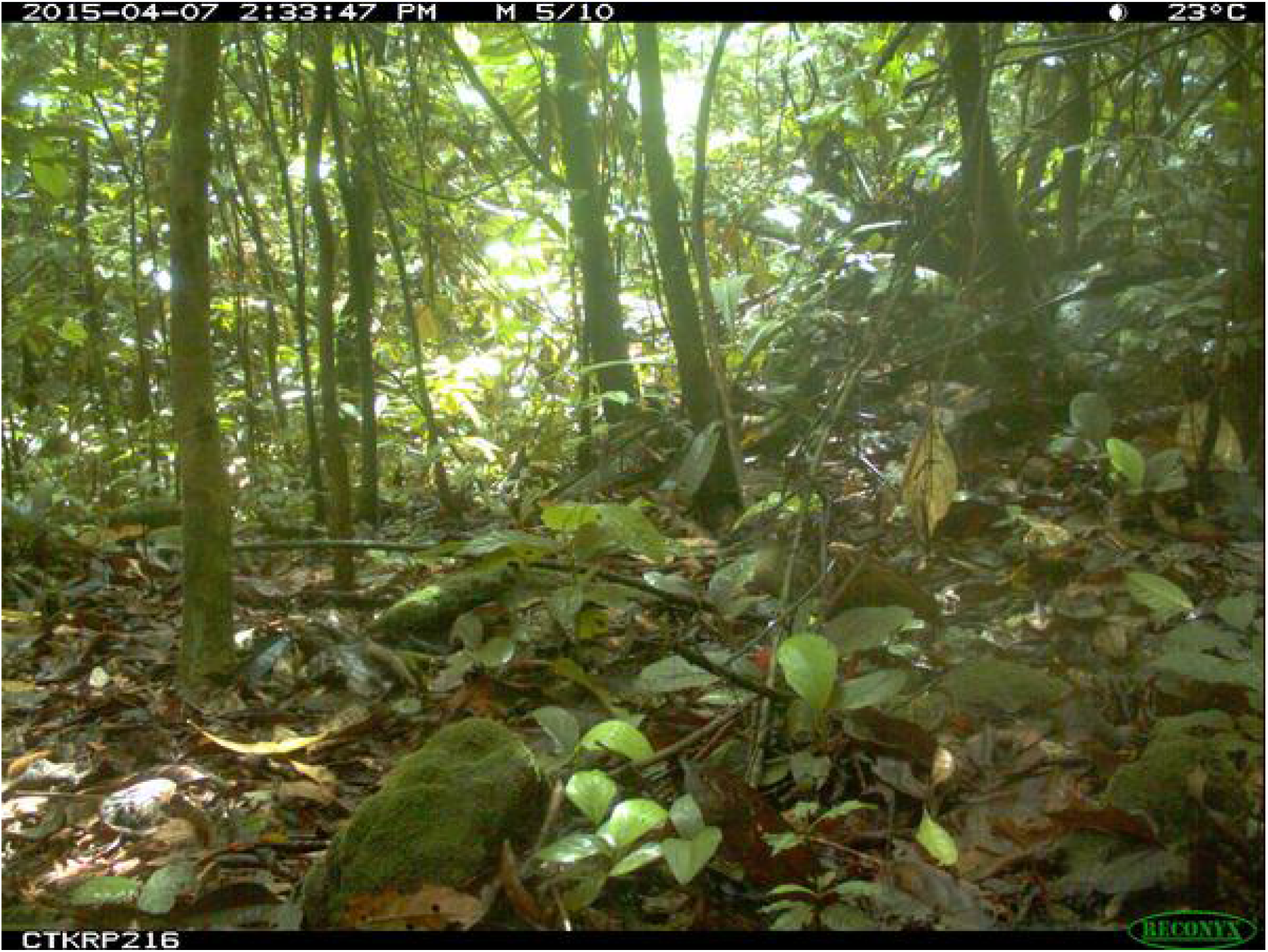
An image correctly classified as nkulengu rail by our machine learning model but marked as blank by an expert. The bird is visible slightly right of center. The dark beak is pointing left and most of the body is hidden behind branches and leaves. A section of its characteristic red legs is visible between the leaves. The model used features from the beak and head region to identify the bird (see Figure S9).

Thresholding improved the overall performance of the model and its performance for individual species. In our tests we ‘discarded’ labels with low confidence but these data could equally be classified manually if sample sizes were small. It is important to note, however, that this additional effort to manually label low confidence images would not have improved inference in our example ecological analyses, with the exception of chimpanzee occupancy estimates. Chimpanzee images had the lowest measure of precision among the four focal species, which suggests that true detection events were probably missed frequently, resulting in false negatives (Figure S2). Species that were classified with the highest precision and accuracy were either relatively unique in their shape, color and pattern (e.g. African leopard, the ‘Genet’ group) or were well represented in the training data. We recommend that users of our model in Central Africa use a threshold of 70% to accept labels and have created an offline, multi-platform software tool that can label large batches of images or videos, and display simple maps of species presence/absence and species richness (available at https://github.com/Appsilon/wildlife-explorer). The software also outputs the labels in a format that can be used for calculating activity patterns or for use in occupancy models. We do not fully automate these analyses at present (in part because of logistical constraints and delays caused by the COVID19 pandemic), but we anticipate these features will be integrated into future releases.

If machine learning models can fully automate labeling of camera trap images, the first question likely to be posed by most ecologists is ‘Should we?’. Camera trap images contain a wealth of information beyond species identity that would be missed using our model such as behavior, demography, individual phenotype and body condition. A trained model is also limited to detecting and classifying the species in the training dataset, and by definition cannot detect new species. Some machine learning models can already classify behavior (Norouzzadeh et al., 2018) and other future models will achieve this and much more. In our opinion fully automated labels can and should be used in ecological analyses, but only after validation (and re-validation) from an ecological perspective, and to answer clearly defined questions. Each use-case will also differ in the benefits that can be gained from fully automated analysis. A conservation manager with tens of thousands of images collected on a rolling basis might accept a trade-off between increased speed of data analysis and having to discard images with uncertain labels, but a scientist testing hypotheses for peer-reviewed publication might prefer to view all of the images manually. We recommend that in all cases models should be validated on a continual basis using sub-sampled data to detect potentially new or hidden biases. Model accuracy could change if field protocols or environmental conditions change in unexpected ways (e.g. heavy snowfall in temperate zones). However, during model evaluation we found that expert labels in the training and validation data were also never themselves ‘perfect’, and perhaps high performance machine learning models offer a more consistent means of analyzing camera trap data than manual labeling because biases are predictable and can be quantified explicitly.

Camera traps are commonly used worldwide by conservation practitioners whose normal scope of work might not allow sufficient time for the handling, processing, and analyzing of large quantities of digital data. The authors personally know of several large camera trap databases that have not been analyzed years after data collection ended, often because of a lack of resources or technical expertise. New web-based platforms for ecological data are seeking to address this problem by allowing users to upload data to the cloud where it is stored and analyzed using machine learning models (Aide et al., 2013; Ahumada et al., 2020), but a lack of fast internet access can be a barrier to using such platforms and our offline application can fill this important gap. The next generation of camera traps will also have embedded machine learning models following the current rise in edge-computing technology. Together, edge and cloud computing will open the door to national and international real-time ecological forecasting at unprecedented spatial and temporal scales. We anticipate that the model, software and validation workflow presented here could revolutionize how camera trap data are processed and analyzed, and conclude that high performance machine learning models can be used for fully automated labeling of camera trap data for ecological analyses.

## Acknowledgments

RW was funded by the EU 11eme FED ECOFAC6 program grant to the National Parks Agency of Gabon. Appsilon Data Science funded the machine learning model and software development costs. Cloud computing costs were funded by a Google Cloud Education Grant awarded to KA. Camera trap data from co-authors KB and CKO were kindly made available by the Tropical Ecology Assessment and Monitoring Network (now https://wildlifeinsights.org).

## Author contributions

R.C.W, J.S., M.R., A.F.K.P., P.H., C.O., R.P., H.R., K.A. T.B. designed research; R.C.W., J.S., M.R. performed research; R.C.W., J.S. analyzed data; R.C.W., J.A.W, L.B., K.B., A.C., D.L., B.M., C.K.O., C.O. collected data; R.W., J.S., J.A.Z., T.B., A.F.K.P, M.R., L.B., K.B., S.B., A.C., P.H., D.L., C.O., H.R., K.A. wrote the paper.

## Data and code availability statement

All data and code will be publicly archived in the University of Stirling’s publicly accessible data repositroy and given a unique DOI on acceptance. Code for the machine learning model is available for review online at https://github.com/Appsilon/gabon_wildlife_training. Code for the offline application to run the model is available at https://github.com/Appsilon/wildlife-explorer. R code for the ecological analyses are available for review online at https://github.com/rcwhytock/Whytock_and_Swiezewski_et_al_2020/.

## Supplementary Information

**Table S1.** Species taxonomy, label descriptions and justification for species/class groups

| Species class | Scientific name | Justification |
| --- | --- | --- |
| Civet_African_Palm | *Nandinia binotata* | - |
| Gorilla | *Gorilla gorilla gorilla* | - |
| Rail_Nkulengu | *Himantornis haematopus* | - |
| Guineafowl_Crested | *Guttera pucherani* | - |
| Mandrillus | *Mandrillus sphinx* | - |
| Blank |  | No animal or human |
| Buffalo_African | *Cyncerus cafer nanus* | - |
| Bird |  | Any other bird |
| Chevrotain_Water | *Hymenoschus aquaticus* | - |
| Guineafowl_Black | *Agelastes niger* | - |
| Cat_Golden | *Caracal aurata* | - |
| Pangolin |  | Identifies any pangolin but trained mainly on *Smutsia gigantea* |
| Duiker_Yellow_Backed | *Cephalophus silvicultor* | - |
| Human | *Homo sapiens* | - |
| Chimpanzee | *Pan troglodytes* | - |
| Monkey |  | Any guenon, colobus or mangabey |
| Mongoose |  | Marsh mongoose *Atilax paludinosus* or long-nosed mongoose *Herpestes naso* |
| Rat_Giant | *Cricetomys emini* | - |
| Duiker_Red | *Cephalophus* sp. | Any of the red *Cephalophus* sp. duikers |
| Duiker_Blue | *Philantomba monticola* | - |
| Hog_Red_River | *Potamochoerus porcus* | - |
| Squirrel |  | Any squirrel but most training data are *Protoxerus stangeri* |
| Leopard_African | *Panthera pardus* | - |
| Elephant_African | *Loxodonta cyclotis* | - |
| Porcupine_Brush_Tailed | *Atherurus africanus* | - |
| Genet | *Genetta* sp. | Most training data are *Genetta servalina* |
| Mongoose_Black_Footed | *Bdeogale nigripes* | - |

**Table S2.** Measures of precision, accuracy and prevalence (%s) for the 27 species/groups (see Table S1 for further details on species groups) in the out-of-sample test data.

| **Species class** | **Precision** | **Recall** | **F1** | **Prevalence** | **Balanced Accuracy** |
| --- | --- | --- | --- | --- | --- |
| Bird | 6.4 | 35.6 | 10.9 | 0.3 | 67 |
| Blank | 96.3 | 31.2 | 47.1 | 13.1 | 65.5 |
| Buffalo_African | 90.6 | 43.3 | 58.6 | 1.6 | 71.6 |
| Cat_Golden | 86.5 | 68.1 | 76.2 | 1.1 | 84 |
| Chevrotain_Water | 96.2 | 37.3 | 53.8 | 0.6 | 68.7 |
| Chimpanzee | 65.3 | 74.5 | 69.6 | 2.4 | 86.8 |
| Civet_African_Palm | 9.1 | 100 | 16.7 | < 0.1 | 100 |
| Duiker_Blue | 73.1 | 91.3 | 81.2 | 14.9 | 92.7 |
| Duiker_Red | 87.5 | 91.8 | 89.6 | 26 | 93.6 |
| Duiker_Yellow_Backed | 88.7 | 71.2 | 79 | 2.9 | 85.5 |
| Elephant_African | 83 | 95 | 88.6 | 15.9 | 95.6 |
| Genet | 87.3 | 93.8 | 90.4 | 0.7 | 96.8 |
| Gorilla | 50 | 15.7 | 23.9 | 0.8 | 57.8 |
| Guineafowl_Black | 22.1 | 60.4 | 32.3 | 0.2 | 80 |
| Guineafowl_Crested | 100 | 16.1 | 27.8 | 0.1 | 58.1 |
| Hog_Red_River | 89.9 | 89.6 | 89.8 | 5.9 | 94.5 |
| Human | 51.5 | 79.9 | 62.6 | 3.6 | 88.6 |
| Leopard_African | 87 | 85.9 | 86.4 | 2 | 92.8 |
| Mandrillus | 73.5 | 26.1 | 38.6 | 2.7 | 62.9 |
| Mongoose | 48.2 | 80.4 | 60.3 | 0.4 | 90 |
| Mongoose_Black_Footed | 72.5 | 64.4 | 68.2 | 0.2 | 82.2 |
| Monkey | 59.9 | 81 | 68.9 | 2.9 | 89.7 |
| Pangolin | 76.7 | 62.2 | 68.7 | 0.2 | 81.1 |
| Porcupine_Brush_Tailed | 86.6 | 83.3 | 84.9 | 0.6 | 91.6 |
| Rail_Nkulengu | 0 | 0 | 0 | 0 | 50 |
| Rat_Giant | 39 | 88.5 | 54.1 | 0.1 | 94.2 |
| Squirrel | 59.9 | 78.2 | 67.8 | 1 | 88.8 |

**Table S3.** Precision, recall, F1 score and prevalence (%s) for the 27 species/groups (see Table S1 for further details on species groups) in the out-of-sample test data at all thresholds used (10 – 90% confidence).

| Species class | Threshold | Precision | Recall | F1 | Prevalence | Balanced Accuracy |
| --- | --- | --- | --- | --- | --- | --- |
| Bird | 10 | 0.064 | 0.356 | 0.109 | 0.003 | 0.670 |
| Blank | 10 | 0.963 | 0.312 | 0.471 | 0.131 | 0.655 |
| Buffalo_African | 10 | 0.906 | 0.433 | 0.586 | 0.016 | 0.716 |
| Cat_Golden | 10 | 0.865 | 0.681 | 0.762 | 0.011 | 0.840 |
| Chevrotain_Water | 10 | 0.962 | 0.373 | 0.538 | 0.006 | 0.687 |
| Chimpanzee | 10 | 0.653 | 0.745 | 0.696 | 0.024 | 0.868 |
| Civet_African_Palm | 10 | 0.091 | 1.000 | 0.167 | 0.000 | 1.000 |
| Duiker_Blue | 10 | 0.731 | 0.913 | 0.812 | 0.149 | 0.927 |
| Duiker_Red | 10 | 0.875 | 0.918 | 0.896 | 0.260 | 0.936 |
| Duiker_Yellow_Backed | 10 | 0.887 | 0.712 | 0.790 | 0.029 | 0.855 |
| Elephant_African | 10 | 0.830 | 0.950 | 0.886 | 0.159 | 0.956 |
| Genet | 10 | 0.873 | 0.938 | 0.904 | 0.007 | 0.968 |
| Gorilla | 10 | 0.500 | 0.157 | 0.239 | 0.008 | 0.578 |
| Guineafowl_Black | 10 | 0.221 | 0.604 | 0.323 | 0.002 | 0.800 |
| Guineafowl_Crested | 10 | 1.000 | 0.161 | 0.278 | 0.001 | 0.581 |
| Hog_Red_River | 10 | 0.899 | 0.896 | 0.898 | 0.059 | 0.945 |
| Human | 10 | 0.515 | 0.799 | 0.626 | 0.036 | 0.886 |
| Leopard_African | 10 | 0.870 | 0.859 | 0.864 | 0.020 | 0.928 |
| Mandrillus | 10 | 0.735 | 0.261 | 0.386 | 0.027 | 0.629 |
| Mongoose | 10 | 0.482 | 0.804 | 0.603 | 0.004 | 0.900 |
| Mongoose_Black_Footed | 10 | 0.725 | 0.644 | 0.682 | 0.002 | 0.822 |
| Monkey | 10 | 0.599 | 0.810 | 0.689 | 0.029 | 0.897 |
| Pangolin | 10 | 0.767 | 0.622 | 0.687 | 0.002 | 0.811 |
| Porcupine_Brush_Tailed | 10 | 0.866 | 0.833 | 0.849 | 0.006 | 0.916 |
| Rail_Nkulengu | 10 | 0.000 | 0.000 | NA | 0.000 | 0.500 |
| Rat_Giant | 10 | 0.390 | 0.885 | 0.541 | 0.001 | 0.942 |
| Squirrel | 10 | 0.599 | 0.782 | 0.678 | 0.010 | 0.888 |
| Bird | 20 | 0.063 | 0.352 | 0.107 | 0.003 | 0.668 |
| Blank | 20 | 0.966 | 0.316 | 0.476 | 0.128 | 0.657 |
| Buffalo_African | 20 | 0.906 | 0.433 | 0.586 | 0.016 | 0.716 |
| Cat_Golden | 20 | 0.865 | 0.688 | 0.767 | 0.011 | 0.844 |
| Chevrotain_Water | 20 | 0.961 | 0.380 | 0.544 | 0.005 | 0.690 |
| Chimpanzee | 20 | 0.659 | 0.745 | 0.699 | 0.024 | 0.868 |
| Civet_African_Palm | 20 | 0.111 | 1.000 | 0.200 | 0.000 | 1.000 |
| Duiker_Blue | 20 | 0.734 | 0.915 | 0.814 | 0.149 | 0.928 |
| Duiker_Red | 20 | 0.877 | 0.919 | 0.898 | 0.261 | 0.937 |
| Duiker_Yellow_Backed | 20 | 0.888 | 0.714 | 0.792 | 0.029 | 0.856 |
| Elephant_African | 20 | 0.830 | 0.951 | 0.886 | 0.160 | 0.957 |
| Genet | 20 | 0.893 | 0.938 | 0.915 | 0.007 | 0.969 |
| Gorilla | 20 | 0.482 | 0.148 | 0.227 | 0.008 | 0.574 |
| Guineafowl_Black | 20 | 0.225 | 0.604 | 0.328 | 0.002 | 0.800 |
| Guineafowl_Crested | 20 | 1.000 | 0.161 | 0.278 | 0.001 | 0.581 |
| Hog_Red_River | 20 | 0.900 | 0.896 | 0.898 | 0.059 | 0.945 |
| Human | 20 | 0.524 | 0.800 | 0.633 | 0.036 | 0.886 |
| Leopard_African | 20 | 0.872 | 0.861 | 0.866 | 0.020 | 0.929 |
| Mandrillus | 20 | 0.735 | 0.263 | 0.388 | 0.027 | 0.630 |
| Mongoose | 20 | 0.516 | 0.804 | 0.628 | 0.004 | 0.900 |
| Mongoose_Black_Footed | 20 | 0.763 | 0.674 | 0.716 | 0.002 | 0.837 |
| Monkey | 20 | 0.601 | 0.814 | 0.691 | 0.029 | 0.899 |
| Pangolin | 20 | 0.821 | 0.622 | 0.708 | 0.002 | 0.811 |
| Porcupine_Brush_Tailed | 20 | 0.866 | 0.853 | 0.859 | 0.005 | 0.926 |
| Rail_Nkulengu | 20 | 0.000 | 0.000 | NA | 0.000 | 0.500 |
| Rat_Giant | 20 | 0.386 | 0.880 | 0.537 | 0.001 | 0.939 |
| Squirrel | 20 | 0.608 | 0.782 | 0.684 | 0.010 | 0.888 |
| Bird | 30 | 0.065 | 0.377 | 0.111 | 0.003 | 0.681 |
| Blank | 30 | 0.968 | 0.329 | 0.491 | 0.112 | 0.664 |
| Buffalo_African | 30 | 0.911 | 0.446 | 0.599 | 0.016 | 0.722 |
| Cat_Golden | 30 | 0.885 | 0.708 | 0.787 | 0.011 | 0.853 |
| Chevrotain_Water | 30 | 0.980 | 0.403 | 0.571 | 0.005 | 0.702 |
| Chimpanzee | 30 | 0.683 | 0.762 | 0.720 | 0.024 | 0.876 |
| Civet_African_Palm | 30 | 0.200 | 1.000 | 0.333 | 0.000 | 1.000 |
| Duiker_Blue | 30 | 0.754 | 0.921 | 0.829 | 0.152 | 0.934 |
| Duiker_Red | 30 | 0.888 | 0.922 | 0.905 | 0.268 | 0.940 |
| Duiker_Yellow_Backed | 30 | 0.909 | 0.726 | 0.807 | 0.029 | 0.862 |
| Elephant_African | 30 | 0.840 | 0.955 | 0.894 | 0.164 | 0.960 |
| Genet | 30 | 0.904 | 0.950 | 0.926 | 0.007 | 0.974 |
| Gorilla | 30 | 0.519 | 0.153 | 0.237 | 0.008 | 0.576 |
| Guineafowl_Black | 30 | 0.283 | 0.604 | 0.386 | 0.002 | 0.800 |
| Guineafowl_Crested | 30 | 1.000 | 0.161 | 0.278 | 0.001 | 0.581 |
| Hog_Red_River | 30 | 0.911 | 0.902 | 0.907 | 0.061 | 0.948 |
| Human | 30 | 0.551 | 0.802 | 0.653 | 0.037 | 0.888 |
| Leopard_African | 30 | 0.887 | 0.872 | 0.879 | 0.020 | 0.935 |
| Mandrillus | 30 | 0.764 | 0.266 | 0.395 | 0.026 | 0.632 |
| Mongoose | 30 | 0.599 | 0.828 | 0.695 | 0.004 | 0.913 |
| Mongoose_Black_Footed | 30 | 0.763 | 0.829 | 0.795 | 0.002 | 0.914 |
| Monkey | 30 | 0.612 | 0.831 | 0.705 | 0.029 | 0.908 |
| Pangolin | 30 | 0.815 | 0.667 | 0.733 | 0.001 | 0.833 |
| Porcupine_Brush_Tailed | 30 | 0.858 | 0.858 | 0.858 | 0.005 | 0.929 |
| Rail_Nkulengu | 30 | 0.000 | 0.000 | NA | 0.000 | 0.500 |
| Rat_Giant | 30 | 0.449 | 0.917 | 0.603 | 0.001 | 0.958 |
| Squirrel | 30 | 0.645 | 0.801 | 0.715 | 0.010 | 0.898 |
| Bird | 40 | 0.078 | 0.423 | 0.132 | 0.002 | 0.706 |
| Blank | 40 | 0.976 | 0.352 | 0.518 | 0.090 | 0.676 |
| Buffalo_African | 40 | 0.929 | 0.473 | 0.627 | 0.015 | 0.736 |
| Cat_Golden | 40 | 0.905 | 0.722 | 0.803 | 0.011 | 0.860 |
| Chevrotain_Water | 40 | 0.977 | 0.452 | 0.618 | 0.004 | 0.726 |
| Chimpanzee | 40 | 0.725 | 0.788 | 0.755 | 0.024 | 0.890 |
| Civet_African_Palm | 40 | 0.200 | 1.000 | 0.333 | 0.000 | 1.000 |
| Duiker_Blue | 40 | 0.791 | 0.934 | 0.857 | 0.157 | 0.944 |
| Duiker_Red | 40 | 0.904 | 0.930 | 0.917 | 0.279 | 0.946 |
| Duiker_Yellow_Backed | 40 | 0.924 | 0.751 | 0.829 | 0.030 | 0.875 |
| Elephant_African | 40 | 0.860 | 0.962 | 0.908 | 0.170 | 0.965 |
| Genet | 40 | 0.921 | 0.950 | 0.935 | 0.007 | 0.975 |
| Gorilla | 40 | 0.528 | 0.119 | 0.195 | 0.007 | 0.559 |
| Guineafowl_Black | 40 | 0.375 | 0.600 | 0.462 | 0.002 | 0.799 |
| Guineafowl_Crested | 40 | 1.000 | 0.161 | 0.278 | 0.001 | 0.581 |
| Hog_Red_River | 40 | 0.930 | 0.911 | 0.920 | 0.063 | 0.953 |
| Human | 40 | 0.593 | 0.811 | 0.685 | 0.038 | 0.895 |
| Leopard_African | 40 | 0.897 | 0.903 | 0.900 | 0.020 | 0.951 |
| Mandrillus | 40 | 0.795 | 0.288 | 0.422 | 0.025 | 0.643 |
| Mongoose | 40 | 0.704 | 0.835 | 0.764 | 0.004 | 0.917 |
| Mongoose_Black_Footed | 40 | 0.800 | 0.903 | 0.848 | 0.001 | 0.951 |
| Monkey | 40 | 0.632 | 0.856 | 0.727 | 0.029 | 0.920 |
| Pangolin | 40 | 0.880 | 0.733 | 0.800 | 0.001 | 0.867 |
| Porcupine_Brush_Tailed | 40 | 0.888 | 0.904 | 0.896 | 0.005 | 0.951 |
| Rail_Nkulengu | 40 | 0.000 | 0.000 | NA | 0.000 | 0.500 |
| Rat_Giant | 40 | 0.537 | 0.917 | 0.677 | 0.001 | 0.958 |
| Squirrel | 40 | 0.715 | 0.843 | 0.774 | 0.010 | 0.920 |
| Bird | 50 | 0.084 | 0.450 | 0.142 | 0.002 | 0.720 |
| Blank | 50 | 0.981 | 0.378 | 0.546 | 0.072 | 0.689 |
| Buffalo_African | 50 | 0.938 | 0.503 | 0.655 | 0.014 | 0.751 |
| Cat_Golden | 50 | 0.923 | 0.741 | 0.822 | 0.011 | 0.870 |
| Chevrotain_Water | 50 | 0.974 | 0.500 | 0.661 | 0.004 | 0.750 |
| Chimpanzee | 50 | 0.753 | 0.824 | 0.787 | 0.024 | 0.909 |
| Civet_African_Palm | 50 | 0.500 | 1.000 | 0.667 | 0.000 | 1.000 |
| Duiker_Blue | 50 | 0.825 | 0.945 | 0.881 | 0.162 | 0.953 |
| Duiker_Red | 50 | 0.920 | 0.939 | 0.930 | 0.288 | 0.953 |
| Duiker_Yellow_Backed | 50 | 0.942 | 0.773 | 0.849 | 0.029 | 0.886 |
| Elephant_African | 50 | 0.879 | 0.968 | 0.921 | 0.177 | 0.969 |
| Genet | 50 | 0.932 | 0.962 | 0.947 | 0.008 | 0.981 |
| Gorilla | 50 | 0.583 | 0.107 | 0.181 | 0.006 | 0.553 |
| Guineafowl_Black | 50 | 0.492 | 0.638 | 0.556 | 0.002 | 0.818 |
| Guineafowl_Crested | 50 | 1.000 | 0.138 | 0.242 | 0.001 | 0.569 |
| Hog_Red_River | 50 | 0.946 | 0.922 | 0.934 | 0.064 | 0.959 |
| Human | 50 | 0.646 | 0.827 | 0.725 | 0.039 | 0.904 |
| Leopard_African | 50 | 0.902 | 0.921 | 0.912 | 0.021 | 0.959 |
| Mandrillus | 50 | 0.816 | 0.292 | 0.430 | 0.023 | 0.645 |
| Mongoose | 50 | 0.775 | 0.868 | 0.819 | 0.004 | 0.934 |
| Mongoose_Black_Footed | 50 | 0.903 | 0.933 | 0.918 | 0.001 | 0.967 |
| Monkey | 50 | 0.654 | 0.879 | 0.750 | 0.029 | 0.932 |
| Pangolin | 50 | 0.913 | 0.724 | 0.808 | 0.001 | 0.862 |
| Porcupine_Brush_Tailed | 50 | 0.902 | 0.953 | 0.927 | 0.005 | 0.976 |
| Rail_Nkulengu | 50 | 0.000 | 0.000 | NA | 0.000 | 0.500 |
| Rat_Giant | 50 | 0.625 | 0.909 | 0.741 | 0.001 | 0.954 |
| Squirrel | 50 | 0.758 | 0.888 | 0.818 | 0.010 | 0.943 |
| Bird | 60 | 0.103 | 0.552 | 0.174 | 0.002 | 0.772 |
| Blank | 60 | 0.985 | 0.399 | 0.568 | 0.052 | 0.699 |
| Buffalo_African | 60 | 0.957 | 0.545 | 0.694 | 0.013 | 0.772 |
| Cat_Golden | 60 | 0.935 | 0.768 | 0.844 | 0.011 | 0.884 |
| Chevrotain_Water | 60 | 0.970 | 0.582 | 0.727 | 0.003 | 0.791 |
| Chimpanzee | 60 | 0.809 | 0.848 | 0.828 | 0.023 | 0.921 |
| Civet_African_Palm | 60 | 0.500 | 1.000 | 0.667 | 0.000 | 1.000 |
| Duiker_Blue | 60 | 0.870 | 0.960 | 0.912 | 0.169 | 0.965 |
| Duiker_Red | 60 | 0.943 | 0.952 | 0.948 | 0.298 | 0.964 |
| Duiker_Yellow_Backed | 60 | 0.962 | 0.802 | 0.875 | 0.029 | 0.901 |
| Elephant_African | 60 | 0.897 | 0.975 | 0.934 | 0.186 | 0.975 |
| Genet | 60 | 0.943 | 0.980 | 0.961 | 0.008 | 0.990 |
| Gorilla | 60 | 0.583 | 0.065 | 0.118 | 0.006 | 0.533 |
| Guineafowl_Black | 60 | 0.614 | 0.711 | 0.659 | 0.002 | 0.855 |
| Guineafowl_Crested | 60 | 1.000 | 0.087 | 0.160 | 0.001 | 0.543 |
| Hog_Red_River | 60 | 0.962 | 0.942 | 0.952 | 0.065 | 0.970 |
| Human | 60 | 0.714 | 0.850 | 0.777 | 0.039 | 0.918 |
| Leopard_African | 60 | 0.916 | 0.945 | 0.930 | 0.022 | 0.971 |
| Mandrillus | 60 | 0.809 | 0.276 | 0.411 | 0.021 | 0.637 |
| Mongoose | 60 | 0.794 | 0.895 | 0.842 | 0.004 | 0.947 |
| Mongoose_Black_Footed | 60 | 0.900 | 1.000 | 0.947 | 0.001 | 1.000 |
| Monkey | 60 | 0.680 | 0.898 | 0.774 | 0.029 | 0.942 |
| Pangolin | 60 | 0.905 | 0.731 | 0.809 | 0.001 | 0.865 |
| Porcupine_Brush_Tailed | 60 | 0.923 | 0.970 | 0.946 | 0.005 | 0.985 |
| Rail_Nkulengu | 60 | 0.000 | 0.000 | NA | 0.000 | 0.500 |
| Rat_Giant | 60 | 0.633 | 0.950 | 0.760 | 0.001 | 0.975 |
| Squirrel | 60 | 0.810 | 0.938 | 0.869 | 0.009 | 0.968 |
| Bird | 70 | 0.112 | 0.600 | 0.189 | 0.001 | 0.797 |
| Blank | 70 | 0.981 | 0.403 | 0.571 | 0.036 | 0.701 |
| Buffalo_African | 70 | 0.975 | 0.557 | 0.709 | 0.012 | 0.778 |
| Cat_Golden | 70 | 0.960 | 0.780 | 0.861 | 0.010 | 0.890 |
| Chevrotain_Water | 70 | 1.000 | 0.674 | 0.806 | 0.002 | 0.837 |
| Chimpanzee | 70 | 0.835 | 0.884 | 0.859 | 0.022 | 0.940 |
| Civet_African_Palm | 70 | NA | NA | NA | 0.000 | NA |
| Duiker_Blue | 70 | 0.904 | 0.970 | 0.936 | 0.176 | 0.974 |
| Duiker_Red | 70 | 0.959 | 0.965 | 0.962 | 0.308 | 0.973 |
| Duiker_Yellow_Backed | 70 | 0.975 | 0.838 | 0.902 | 0.029 | 0.919 |
| Elephant_African | 70 | 0.919 | 0.984 | 0.951 | 0.193 | 0.982 |
| Genet | 70 | 0.953 | 0.993 | 0.972 | 0.008 | 0.996 |
| Gorilla | 70 | 0.000 | 0.000 | NA | 0.004 | 0.500 |
| Guineafowl_Black | 70 | 0.706 | 0.727 | 0.716 | 0.002 | 0.863 |
| Guineafowl_Crested | 70 | 1.000 | 0.053 | 0.100 | 0.001 | 0.526 |
| Hog_Red_River | 70 | 0.970 | 0.957 | 0.963 | 0.065 | 0.977 |
| Human | 70 | 0.784 | 0.874 | 0.826 | 0.040 | 0.932 |
| Leopard_African | 70 | 0.928 | 0.960 | 0.944 | 0.022 | 0.979 |
| Mandrillus | 70 | 0.839 | 0.290 | 0.431 | 0.018 | 0.645 |
| Mongoose | 70 | 0.835 | 0.910 | 0.871 | 0.004 | 0.955 |
| Mongoose_Black_Footed | 70 | 0.929 | 1.000 | 0.963 | 0.001 | 1.000 |
| Monkey | 70 | 0.707 | 0.920 | 0.800 | 0.029 | 0.954 |
| Pangolin | 70 | 0.941 | 0.800 | 0.865 | 0.001 | 0.900 |
| Porcupine_Brush_Tailed | 70 | 0.939 | 0.989 | 0.963 | 0.005 | 0.994 |
| Rail_Nkulengu | 70 | 0.000 | 0.000 | NA | 0.000 | 0.500 |
| Rat_Giant | 70 | 0.682 | 0.938 | 0.789 | 0.001 | 0.969 |
| Squirrel | 70 | 0.859 | 0.958 | 0.906 | 0.009 | 0.978 |
| Bird | 80 | 0.151 | 0.786 | 0.253 | 0.001 | 0.891 |
| Blank | 80 | 0.986 | 0.363 | 0.530 | 0.023 | 0.681 |
| Buffalo_African | 80 | 1.000 | 0.596 | 0.747 | 0.010 | 0.798 |
| Cat_Golden | 80 | 0.962 | 0.839 | 0.896 | 0.009 | 0.919 |
| Chevrotain_Water | 80 | 1.000 | 0.750 | 0.857 | 0.002 | 0.875 |
| Chimpanzee | 80 | 0.875 | 0.919 | 0.897 | 0.019 | 0.958 |
| Civet_African_Palm | 80 | NA | NA | NA | 0.000 | NA |
| Duiker_Blue | 80 | 0.932 | 0.981 | 0.956 | 0.183 | 0.982 |
| Duiker_Red | 80 | 0.973 | 0.975 | 0.974 | 0.315 | 0.981 |
| Duiker_Yellow_Backed | 80 | 0.988 | 0.885 | 0.934 | 0.028 | 0.942 |
| Elephant_African | 80 | 0.940 | 0.991 | 0.965 | 0.203 | 0.987 |
| Genet | 80 | 0.949 | 1.000 | 0.974 | 0.008 | 1.000 |
| Gorilla | 80 | 0.000 | 0.000 | NA | 0.003 | 0.500 |
| Guineafowl_Black | 80 | 0.769 | 0.690 | 0.727 | 0.002 | 0.845 |
| Guineafowl_Crested | 80 | 1.000 | 0.067 | 0.125 | 0.001 | 0.533 |
| Hog_Red_River | 80 | 0.980 | 0.971 | 0.975 | 0.065 | 0.985 |
| Human | 80 | 0.853 | 0.892 | 0.872 | 0.040 | 0.943 |
| Leopard_African | 80 | 0.952 | 0.974 | 0.963 | 0.023 | 0.987 |
| Mandrillus | 80 | 0.888 | 0.305 | 0.454 | 0.014 | 0.652 |
| Mongoose | 80 | 0.829 | 0.944 | 0.883 | 0.004 | 0.972 |
| Mongoose_Black_Footed | 80 | 1.000 | 1.000 | 1.000 | 0.001 | 1.000 |
| Monkey | 80 | 0.756 | 0.928 | 0.833 | 0.029 | 0.959 |
| Pangolin | 80 | 1.000 | 0.824 | 0.903 | 0.001 | 0.912 |
| Porcupine_Brush_Tailed | 80 | 0.935 | 0.989 | 0.961 | 0.005 | 0.994 |
| Rail_Nkulengu | 80 | NA | NA | NA | 0.000 | NA |
| Rat_Giant | 80 | 0.737 | 0.933 | 0.824 | 0.001 | 0.967 |
| Squirrel | 80 | 0.879 | 0.979 | 0.926 | 0.008 | 0.989 |
| Bird | 90 | 0.220 | 0.900 | 0.353 | 0.001 | 0.949 |
| Blank | 90 | 1.000 | 0.320 | 0.484 | 0.011 | 0.660 |
| Buffalo_African | 90 | 1.000 | 0.647 | 0.785 | 0.008 | 0.823 |
| Cat_Golden | 90 | 0.980 | 0.897 | 0.937 | 0.007 | 0.949 |
| Chevrotain_Water | 90 | 1.000 | 0.833 | 0.909 | 0.001 | 0.917 |
| Chimpanzee | 90 | 0.914 | 0.922 | 0.918 | 0.016 | 0.960 |
| Civet_African_Palm | 90 | NA | NA | NA | 0.000 | NA |
| Duiker_Blue | 90 | 0.957 | 0.990 | 0.973 | 0.196 | 0.989 |
| Duiker_Red | 90 | 0.984 | 0.984 | 0.984 | 0.317 | 0.988 |
| Duiker_Yellow_Backed | 90 | 0.994 | 0.912 | 0.952 | 0.026 | 0.956 |
| Elephant_African | 90 | 0.961 | 0.994 | 0.977 | 0.218 | 0.991 |
| Genet | 90 | 0.957 | 1.000 | 0.978 | 0.007 | 1.000 |
| Gorilla | 90 | NA | 0.000 | NA | 0.002 | 0.500 |
| Guineafowl_Black | 90 | 0.864 | 0.826 | 0.844 | 0.002 | 0.913 |
| Guineafowl_Crested | 90 | NA | 0.000 | NA | 0.001 | 0.500 |
| Hog_Red_River | 90 | 0.986 | 0.982 | 0.984 | 0.063 | 0.990 |
| Human | 90 | 0.918 | 0.923 | 0.920 | 0.040 | 0.960 |
| Leopard_African | 90 | 0.973 | 0.979 | 0.976 | 0.025 | 0.989 |
| Mandrillus | 90 | 0.900 | 0.298 | 0.448 | 0.010 | 0.649 |
| Mongoose | 90 | 0.855 | 0.967 | 0.908 | 0.004 | 0.983 |
| Mongoose_Black_Footed | 90 | 1.000 | 1.000 | 1.000 | 0.001 | 1.000 |
| Monkey | 90 | 0.811 | 0.952 | 0.876 | 0.028 | 0.973 |
| Pangolin | 90 | 1.000 | 0.813 | 0.897 | 0.001 | 0.906 |
| Porcupine_Brush_Tailed | 90 | 0.934 | 1.000 | 0.966 | 0.005 | 1.000 |
| Rail_Nkulengu | 90 | NA | NA | NA | 0.000 | NA |
| Rat_Giant | 90 | 0.846 | 0.917 | 0.880 | 0.001 | 0.958 |
| Squirrel | 90 | 0.938 | 0.981 | 0.959 | 0.007 | 0.990 |

**Table S4.**
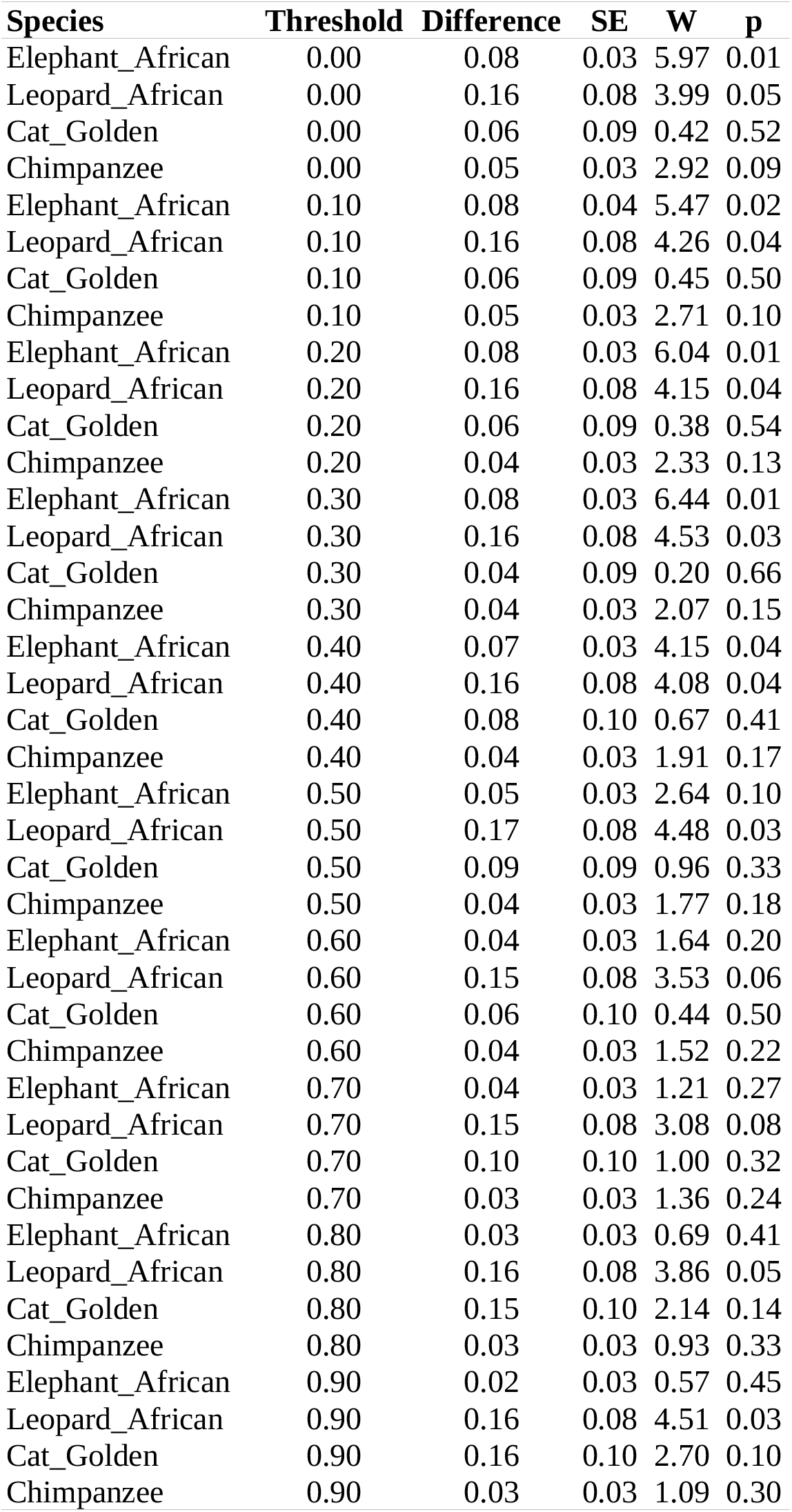
Difference in proportion of day (24 h) active for each species and threshold combination showing standard error (SE), Wald test statistic (W) and p value (p).

**Figure S1.**
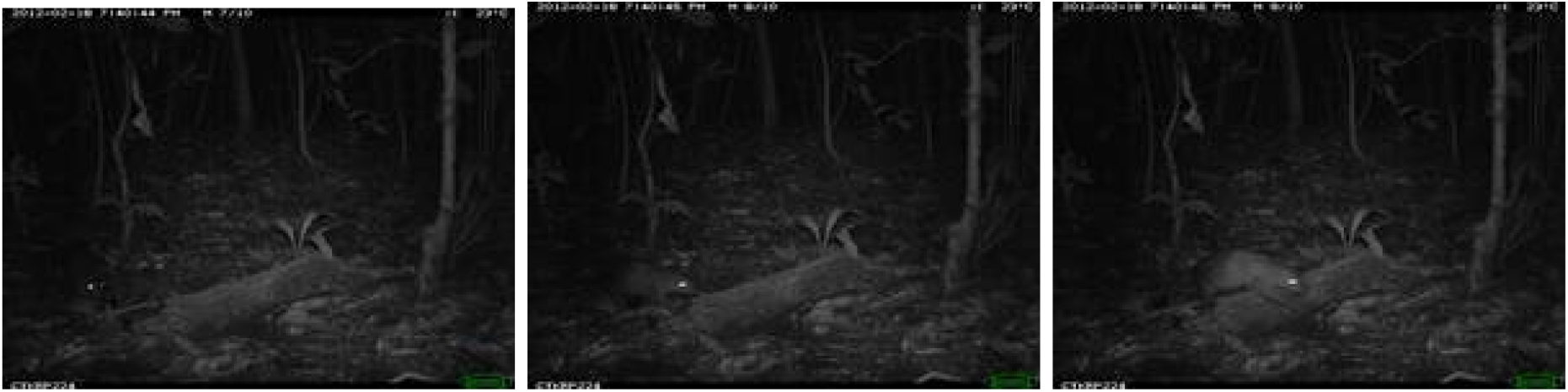
Three example photos taken from a burst of 10 images, showing a porcupine *Atherurus africanus* walking in front of the camera.

**Figure S2.**
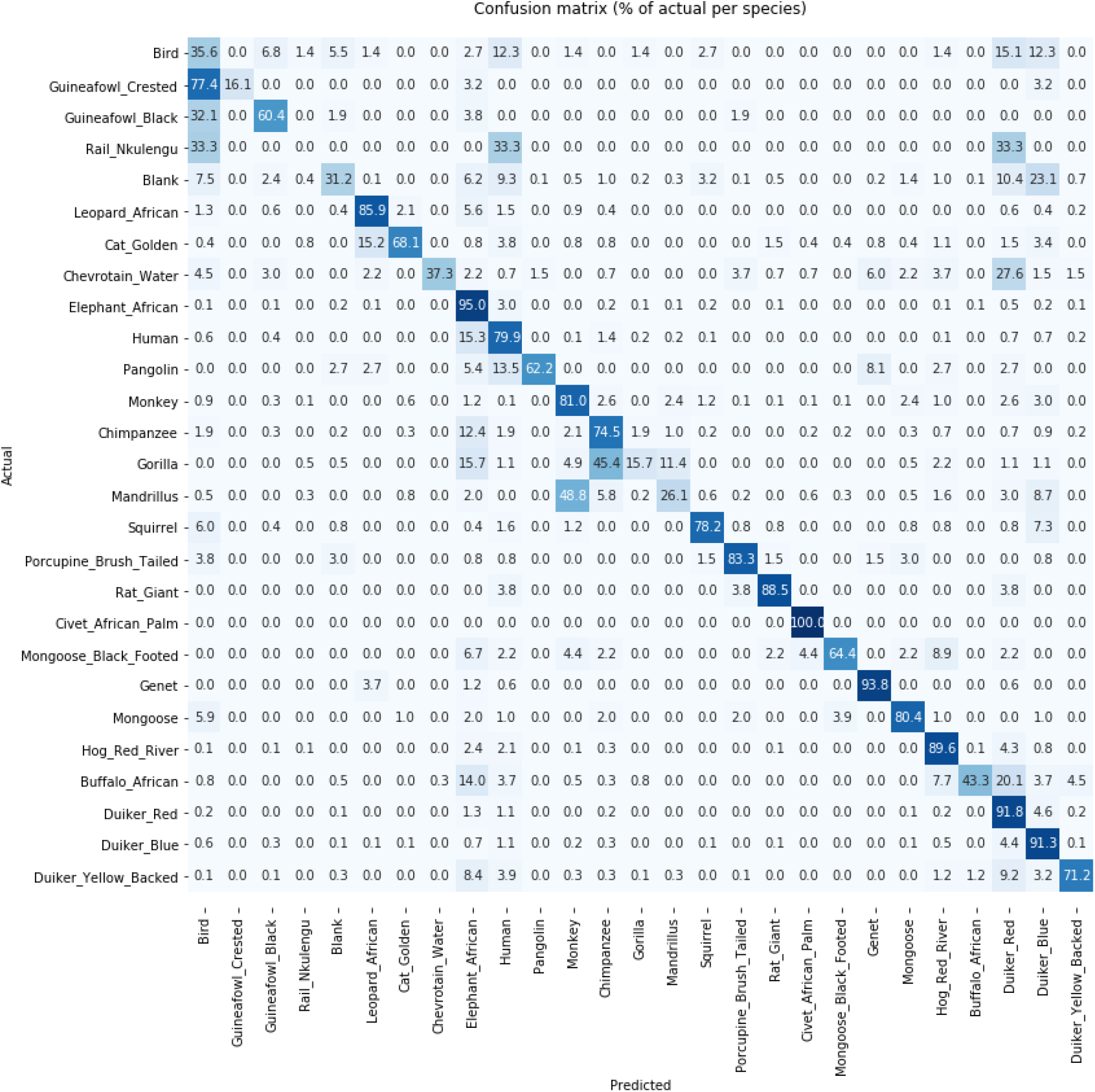
Confusion matrix showing model performance on out of sample test data (each row is normalized independently). Figure S7 shows the confusion matrix with absolute numbers.

**Figure S3.**
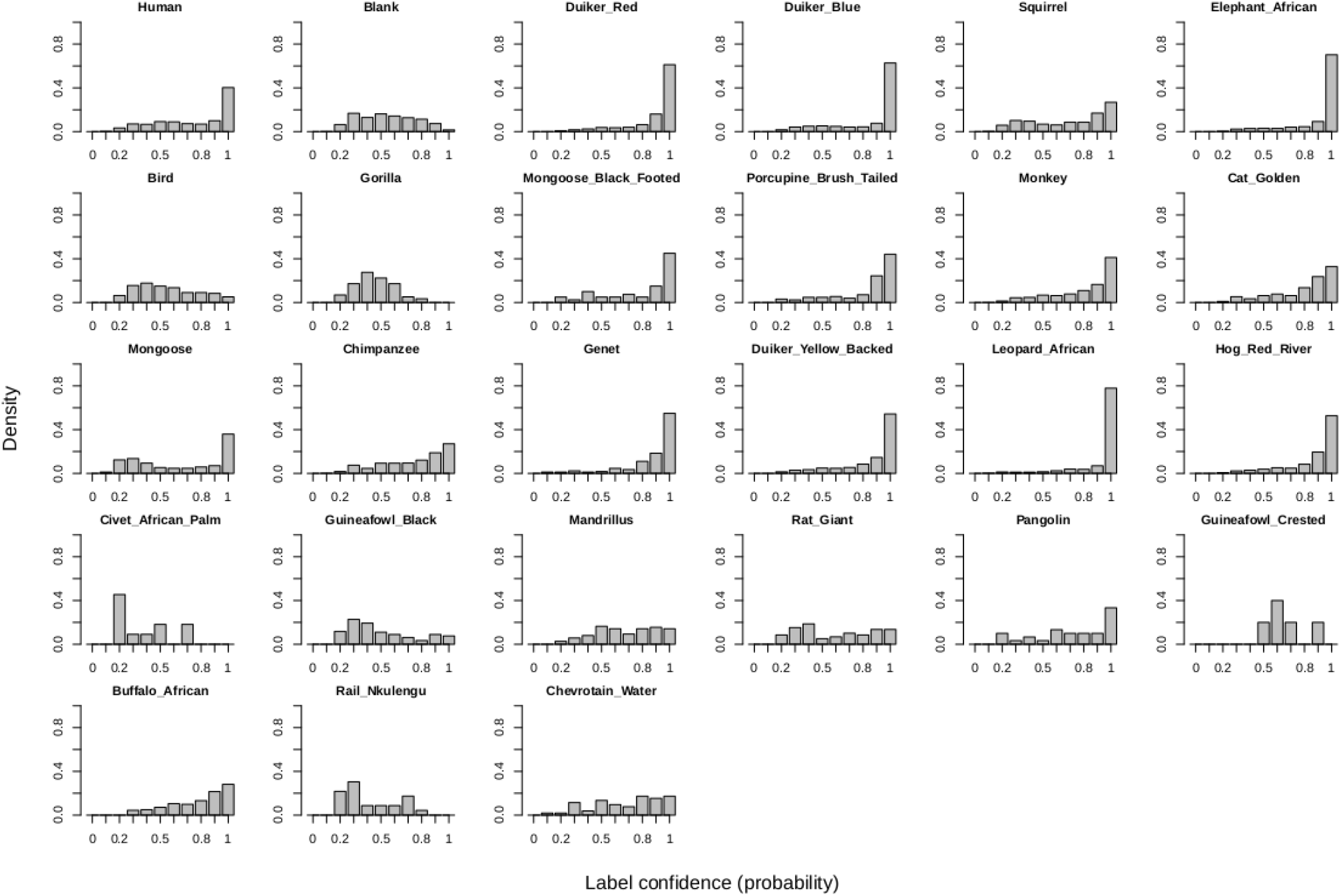
Histograms showing the frequency distribution (normalized density) of label confidence from the machine learning model for the 27 classes in the out-of-sample test data.

**Figure S4.**
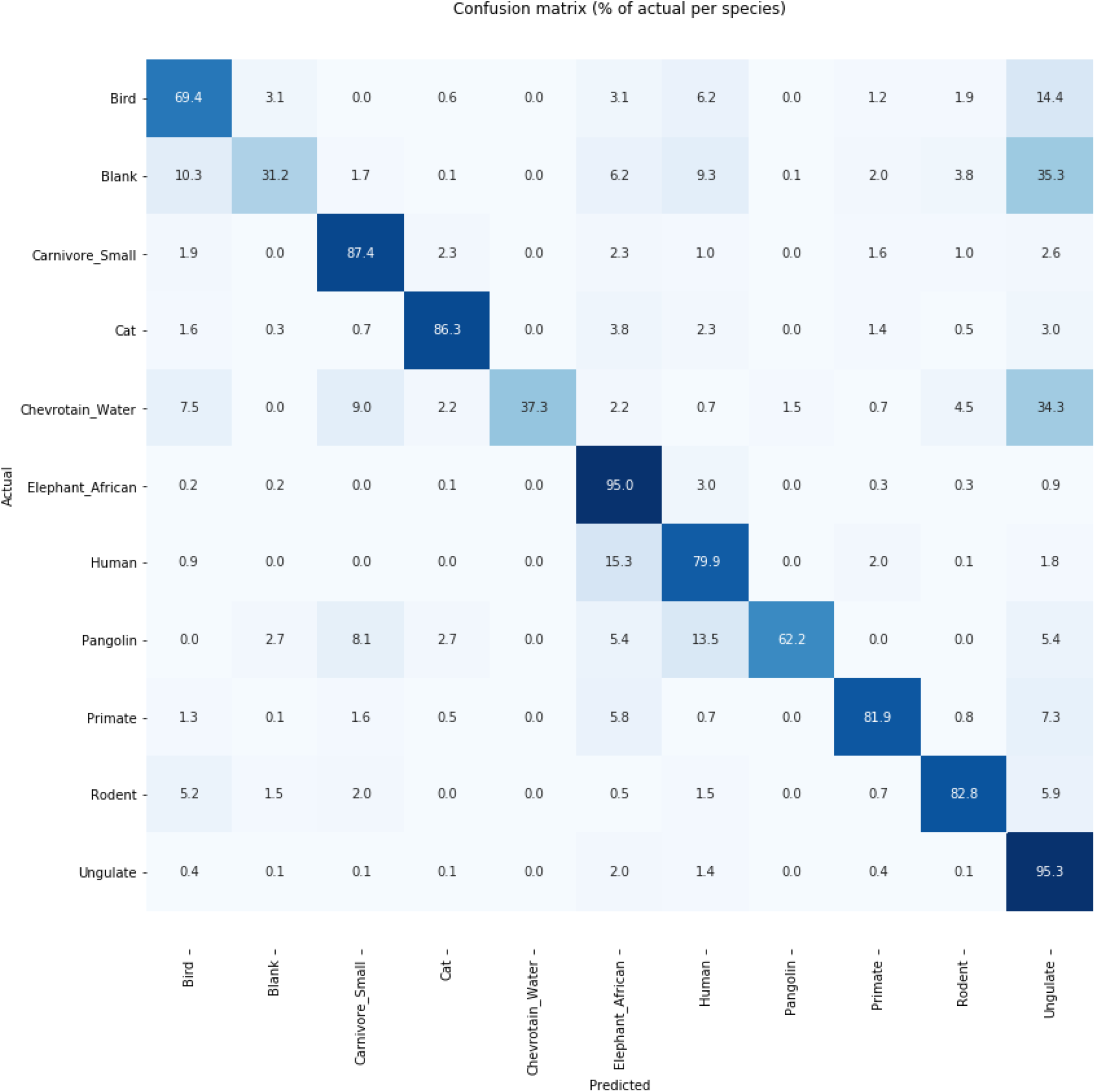
Confusion matrix showing model performance for an aggregated set of 11 classes.

**Figure S5.**
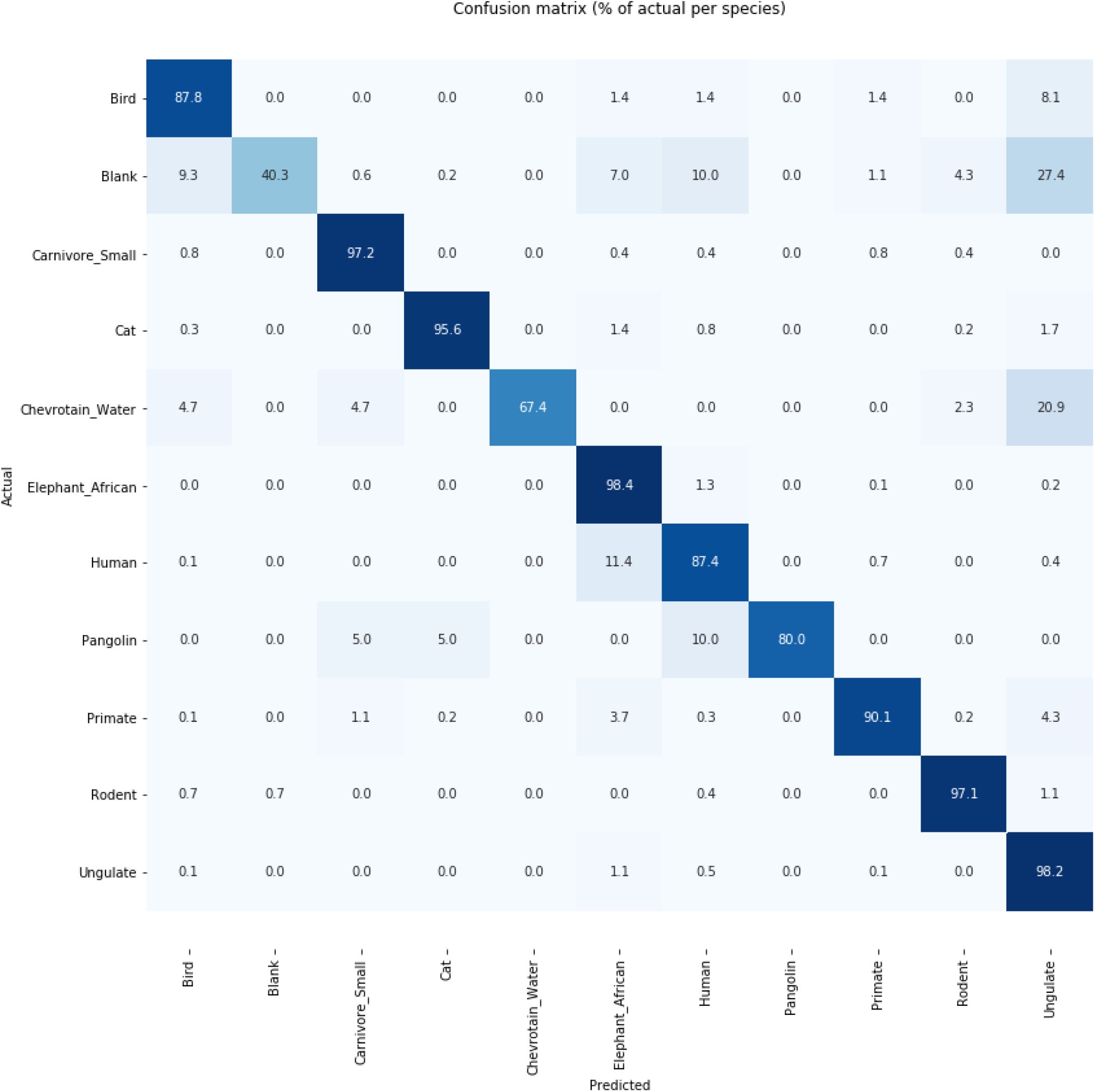
Confusion matrix showing model performance for an aggregated set of 11 classes after removing labels with a predicted confidence < 70%

**Figure S6.**
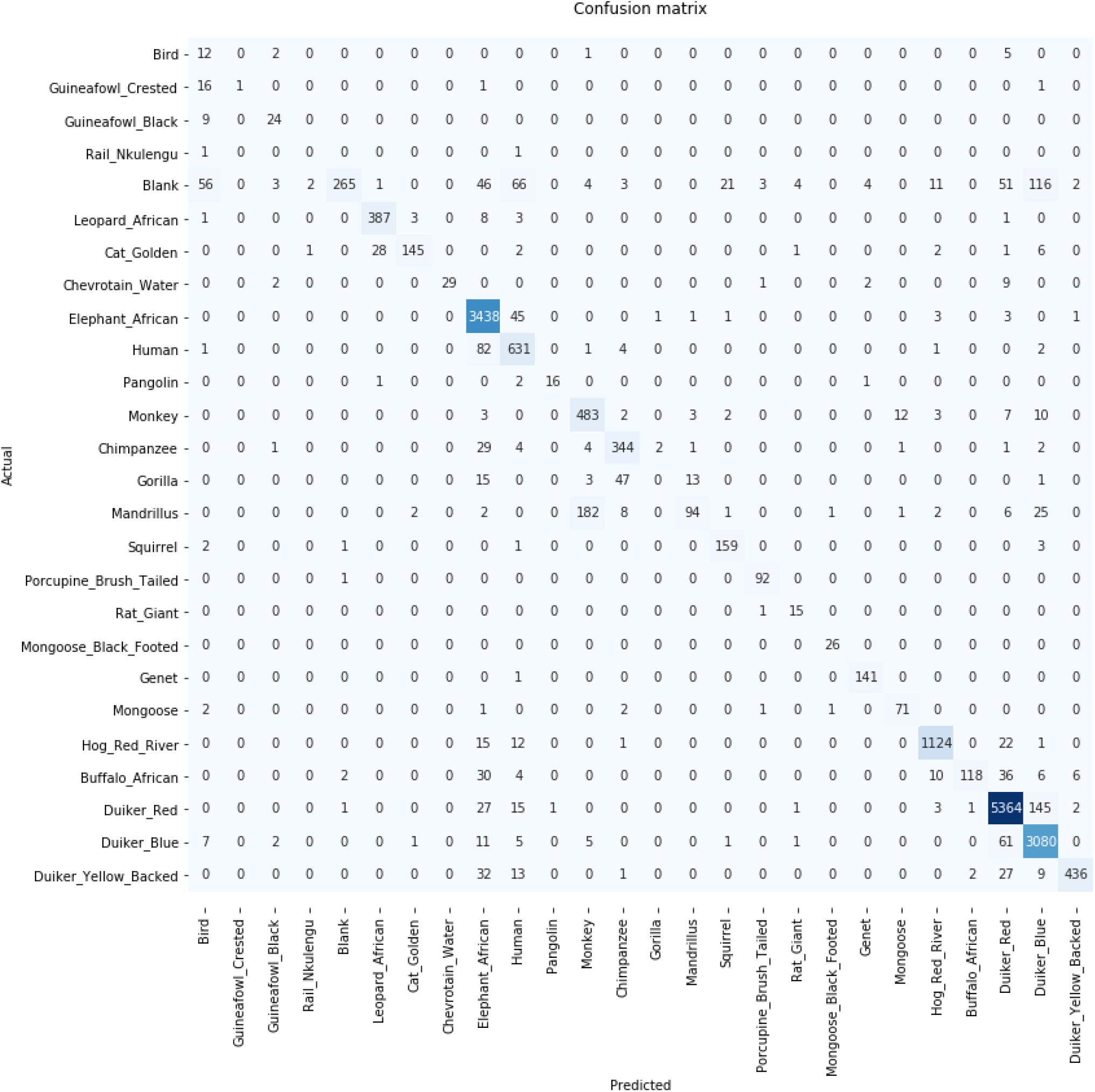
Confusion matrix showing model performance on out of sample test data after excluding labels below a confidence threshold of 70% (with absolute numbers). Figure 3 shows the confusion matrix with each row normalized independently.

**Figure S7.**
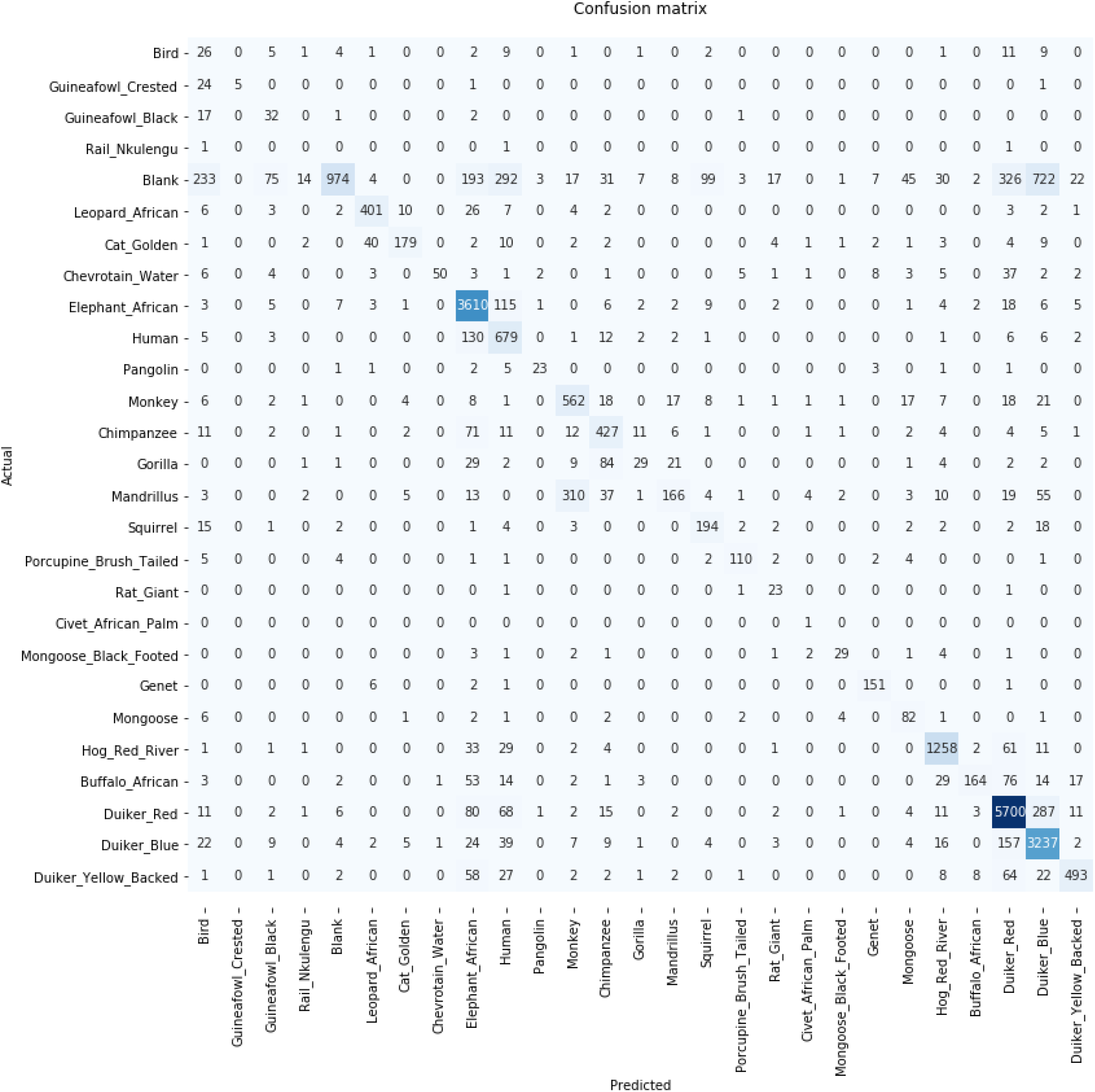
Confusion matrix showing model performance on out of sample test data (absolute numbers). Figure S2 shows the confusion matrix with each row normalized independently.

**Figure S8.**
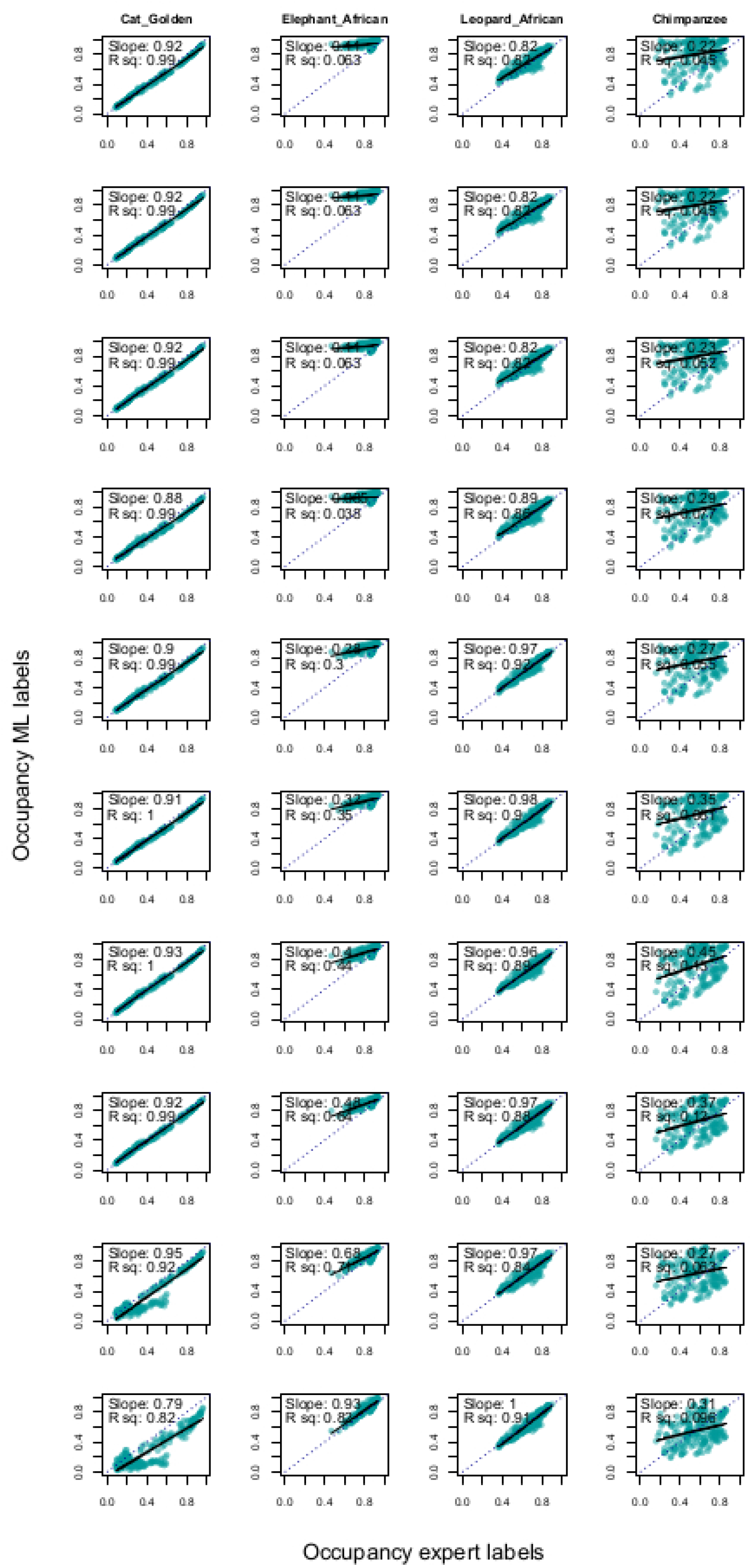
Relationship between estimated occupancy probability for *n* = 227 camera stations (points) from machine learning (ML) labels (y-axis) and expert labels (x-axis) for the four focal species at each threshold (row) from 0 to 90%, in 10% intervals.

**Figure S9.**
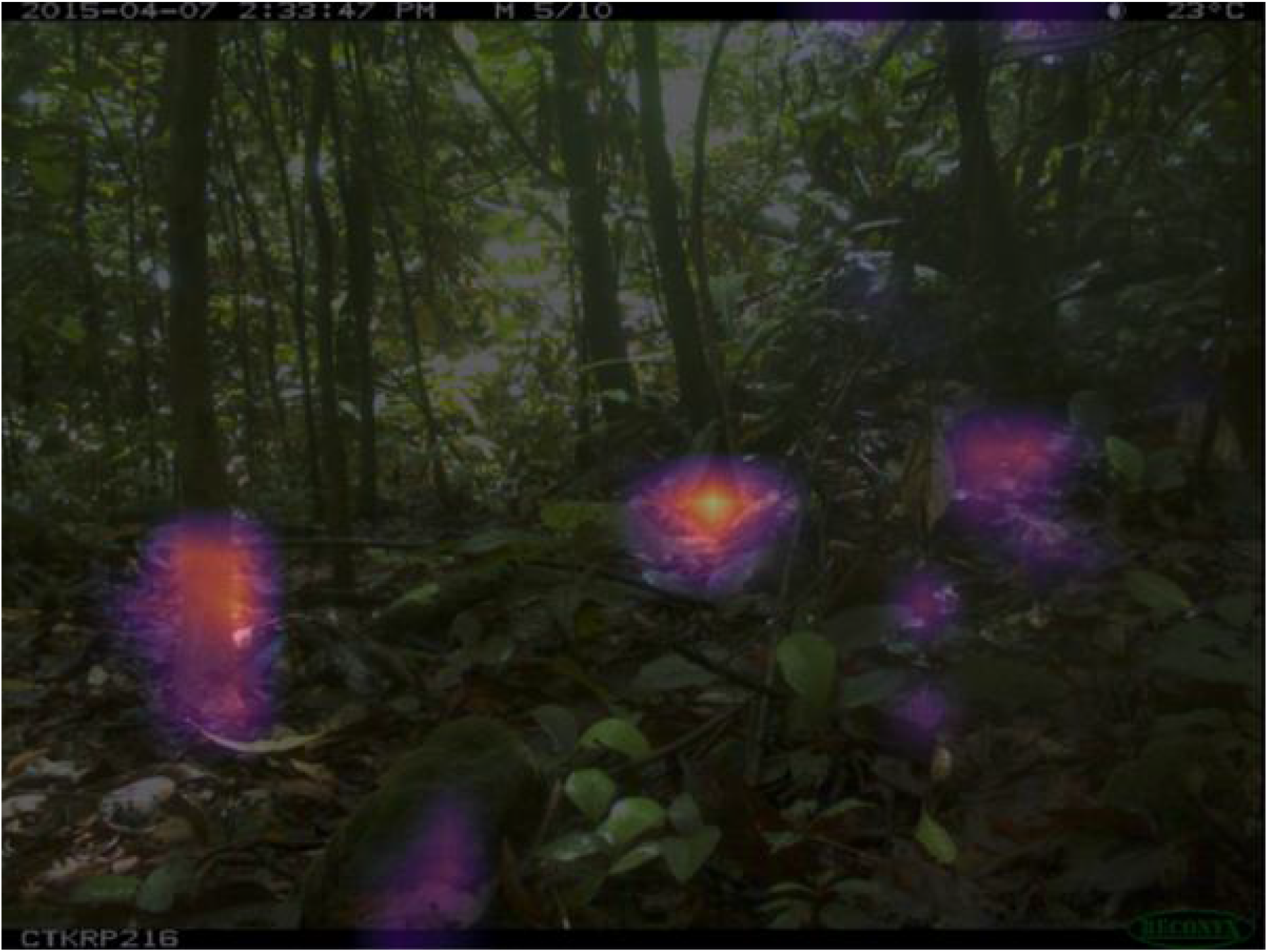
The image from Figure 9 with an added layer illustrating the most important regions of the image for the model when identifying the nkulengu rail. The brightest spot (yellow) near the center of the image encompasses a part of the bird’s beak and head, which apparently were crucial during identification. We used the Grad-CAM (1) technique to create this image.

## Notes

https://github.com/Appsilon/gabon_wildlife_training

https://github.com/rcwhytock/Whytock_and_Swiezewski_et_al_2020/

### Competing Interest Statement

The authors have declared no competing interest.

### Summary of Updates

Updated author Orcid ID. Updated one author name. Revised Abstract and referencing to conform to journal guidelines.

